# *De novo* Design of All-atom Biomolecular Interactions with RFdiffusion3

**DOI:** 10.1101/2025.09.18.676967

**Authors:** Jasper Butcher, Rohith Krishna, Raktim Mitra, Rafael I. Brent, Yanjing Li, Nathaniel Corley, Paul T. Kim, Jonathan Funk, Simon Mathis, Saman Salike, Aiko Muraishi, Helen Eisenach, Tuscan Rock Thompson, Jie Chen, Yuliya Politanska, Enisha Sehgal, Brian Coventry, Odin Zhang, Bo Qiang, Kieran Didi, Max Kazman, Frank DiMaio, David Baker

## Abstract

Deep learning has accelerated protein design, but most existing methods are restricted to generating protein backbone coordinates and often neglect interactions with other biomolecules. We present RFdiffusion3 (RFD3), a diffusion model that generates protein structures in the context of ligands, nucleic acids and other non-protein constellations of atoms. Because all polymer atoms are modeled explicitly, conditioning the model on complex sets of atom-level constraints for enzyme design and other challenges is both simpler and more effective than previous approaches. RFD3 achieves improved performance compared to prior approaches on a range of *in silico* benchmarks with one tenth the computational cost. Finally, we demonstrate the broad applicability of RFD3 by designing and experimentally characterizing DNA binding proteins and cysteine hydrolases. The ability to rapidly generate protein structures guided by complex sets of atom-level constraints in the context of arbitrary non-protein atoms should further expand the range of functions attainable through protein design.

## 1 Main

There has been considerable recent progress in *de novo* design of proteins with new functions using generative deep learning methods [1, 2, 3, 4, 5, 6, 7, 8, 9, 10]. Most methods, such as RFdiffusion (RFD1) [1] and BindCraft [4], represent proteins at the amino-acid level. This granularity empirically suffices for designing protein monomers, assemblies, and protein-binding proteins, but lacks the resolution to design specific side-chain interactions with non-protein atoms such as small molecules and nucleic acids. RFdiffusion2 (RFD2) [3] addressed this challenge by enabling the conditioning of residue-level diffusion on a small number of side chain atoms critical for catalysis or ligand binding; however, because diffusion remains at the residue level, it is incapable of creating additional side chain interactions with ligands or catalytic residues. In structure prediction, AlphaFold3 (AF3) [11] utilized a diffusion process over every individual atom conditioned on rich representations of protein geometry generated by a separate conditioning module. Further work [12, 13, 14]c showed that atomic diffusion processes can generate protein backbones and can be extended to model side chains, but these efforts did not enable modeling of interactions with non-protein components.

We hypothesized that a general solution to the challenge of designing systems of proteins interacting with small molecules, nucleic acids and other non-protein components could be achieved using atom-based diffusion including both backbone and sidechain atoms while conditioning on the non-protein components of the system and the positions of key protein atoms contributing to function. We set out to develop such a model for a wide range of current design challenges (Fig. 1b).

**Figure 1:**
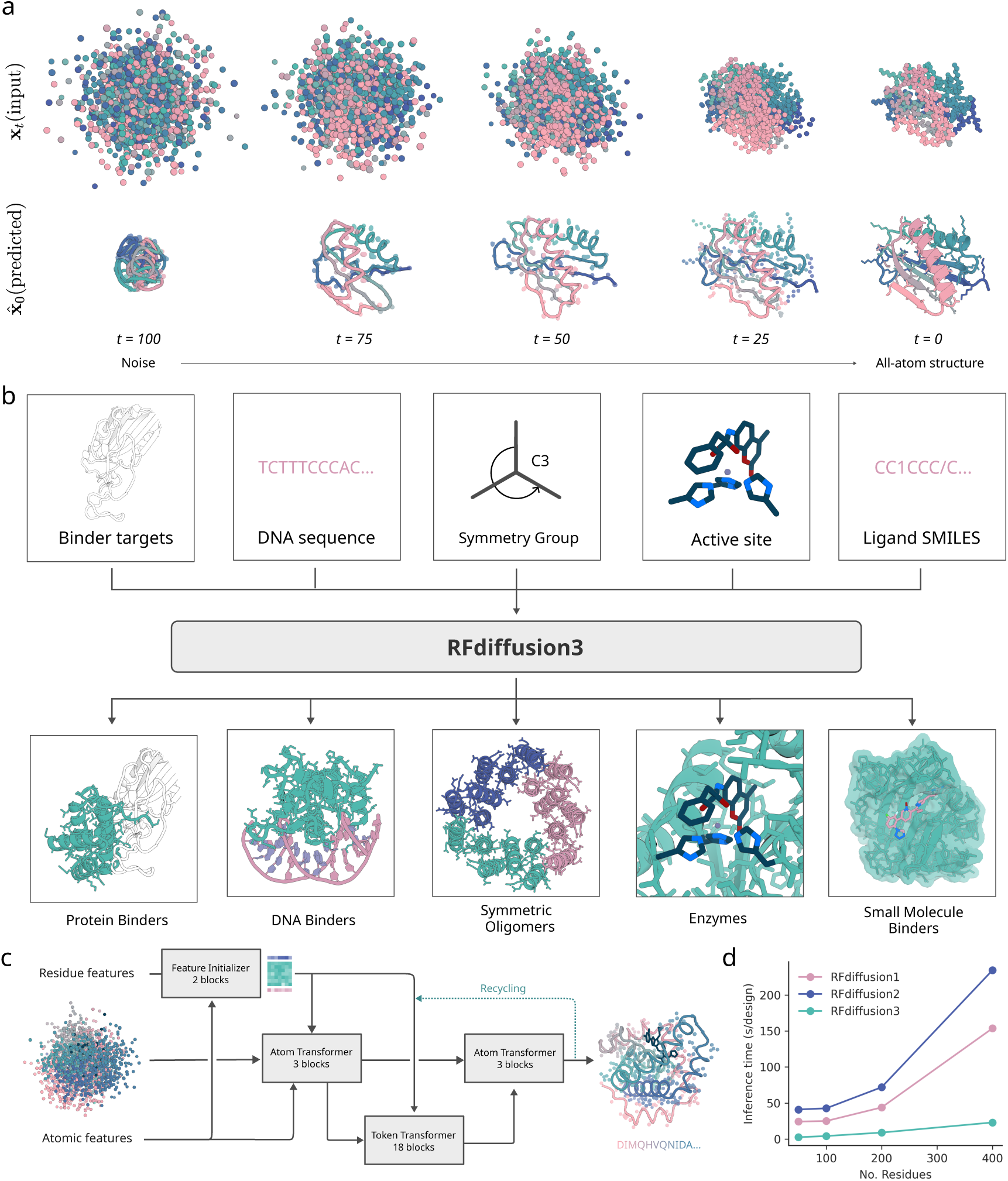
All-atom protein design with RFdiffusion3. **a,** Depiction of 100 step RFD3 diffusion trajectory, predicting both backbone and sidechain atoms. (top) noisy co-ordinates at different timesteps. (bottom) the network prediction at each time-step. **b,** Generation of biomolecular interactions. (top) different conditions can be fed to the model. (bottom) sample output structures for each input condition. **c,** Model architecture. Arrows show information flow, grey boxes indicate modules with learnable parameters. **d,** Inference times for RFdiffusion variants across length scales. RFD3 is an order of magnitude faster than previous versions of RFdiffusion because of its simplified architecture (runtimes are measured on NVIDIA A6000 GPUs).

## 2 RFdiffusion3 architecture and training

Using the AtomWorks framework [15] ^1^, we set out to explore a generative approach in which the fundamental units being diffused are atoms rather than amino acid residues. A first challenge in using such an approach for protein design is how to represent side chains when the sequence is unknown, since amino acids have different numbers of side chain atoms. To address this issue, we treat all residues as having 4 backbone atoms and 10 side chain atoms (the largest canonical sidechain, tryptophan, has 10 atoms) (Figure S1). We supplement the smaller sidechains with additional virtual atoms placed on the C*_β_* [16].

To generate atomic coordinates starting from random initializations, we adopted a transformer-based U-Net architecture, similarly to the AlphaFold3 diffusion module, with three parts: (i) a downsampling module that encodes atomic- and residue-level features including the partially noised structure, (ii) a sparse transformer module that processes token-wise information, and (iii) an upsampling module that modulates the atomic features with the token-wise features and predicts coordinate updates (Fig. 1c) [11, 17]. Joint sampling of backbone and side chain coordinates requires effective coupling between atom-level and residue-level features; we therefore introduce two new architectural elements for feature exchange. First, information transfer is conducted through a series of sparse attention blocks between atomic and token features, only allowing residues and atoms that are geometrically close in the noise to attend to each other. These distance-based updates enable the network to focus on local interactions and prevent overfitting. Second, inspired by Pagnoni et al. [18], we use a cross attention mechanism to up-pool and down-pool between the fine-grained atom representations and coarser-grained token representations. In these updates, the features of each atom are weighted by an attention operation before they are pooled into token features, enabling the token-level transformer to focus on specific atoms within each token. After processing, the token features are read out from the token track using a cross attention by token splitting (Figure S12).

Protein design differs from protein structure prediction in that the input is not an amino acid sequence but instead a set of high-level design conditions (protein length, possible target molecules, etc). Whereas AF3 uses the time- and compute-intensive Pairformer module to extract distance and other information from the input amino acid sequence, for design we reasoned that a much lighter weight module could suffice for processing conditioning information. We thus shrunk the Pairformer from 48 to just 2 layers, greatly reducing computational cost, and ultimately yielding only 168M trainable parameters for the final network (vs. ∼ 350M for AF3). To further improve generation efficiency, we omit computationally expensive triangle multiplicative and triangle attention updates from the network, widely used in prior architectures for protein design [1, 2, 3, 16, 12] and structure prediction [11, 19, 2, 20]. The processed conditioning information is input into the modified diffusion module described above, where most of the compute time is spent.

Design of proteins with new functions generally requires the specification of multiple coarse- and fine-grained constraints. RFD3 extends the conditioning capabilities of RFdiffusion2 (RFD2) [3], which allows coordinate-level constraints such as functional motifs with or without sequence indexing, and coarse-grained constraints such as solvent exposure of ligands. The natively all-atom architecture of RFD3 considerably simplifies the specification of atomic-level constraints, providing precise control over hydrogen bonding, ligand contacts, and interactions with nucleic acids. For binder design and motif scaffolding challenges, the coordinates of the motif or binding partner are provided as exact information within the developing noise cloud, allowing the network to refine the structure in the context of the motif. This approach ensures that the generated proteins respect the precise geometry of the given functional motifs. We used classifier-free guidance, originally developed in the context of image diffusion [21], to improve adherence to the conditioning information. At each denoising step, the network performs a weighted average of two forward passes: one using the conditioning information and one unconditional. We found that this considerably improved the ability of the model to satisfy complex sets of conditions.

Beyond providing target and motif atomic conditioning information, we trained RFD3 to follow several additional types of conditioning. Atomic-resolution modeling has the advantage of enabling direct specification of atoms that should participate in hydrogen bonding interactions with the diffused protein. We trained RFD3 providing in a small fraction of cases information about which atoms are hydrogen bond donors or acceptors. We found that for small molecule binder design, conditioning on hydrogen bond donor or acceptor increases the fraction of such interactions from 26.67% to 32.67%, and this fraction can be further enhanced to 36.67% using classifier-free guidance (Fig. 3d. Similar trends are observed for hydrogen bonds to DNA bases (11% vs 11.3% vs 12.5%). To enable specifying the extent of burial of ligand atoms and their positions relative to the centre-of-mass of the generated protein, we enable users to provide solvent-accessible surface area labels (Fig. 2e) and the position of the generated protein centre of mass relative to a target molecule or motif (Fig. 2f). Similar to RFdiffusion1, providing symmetric noise as input enables generating symmetric structures (Fig. 2g; further *in silico* benchmarking in Fig. S6). Together, these conditioning mechanisms provide considerable control over the generated structures, enabling the design of a wide range of functional proteins.

**Figure 2:**
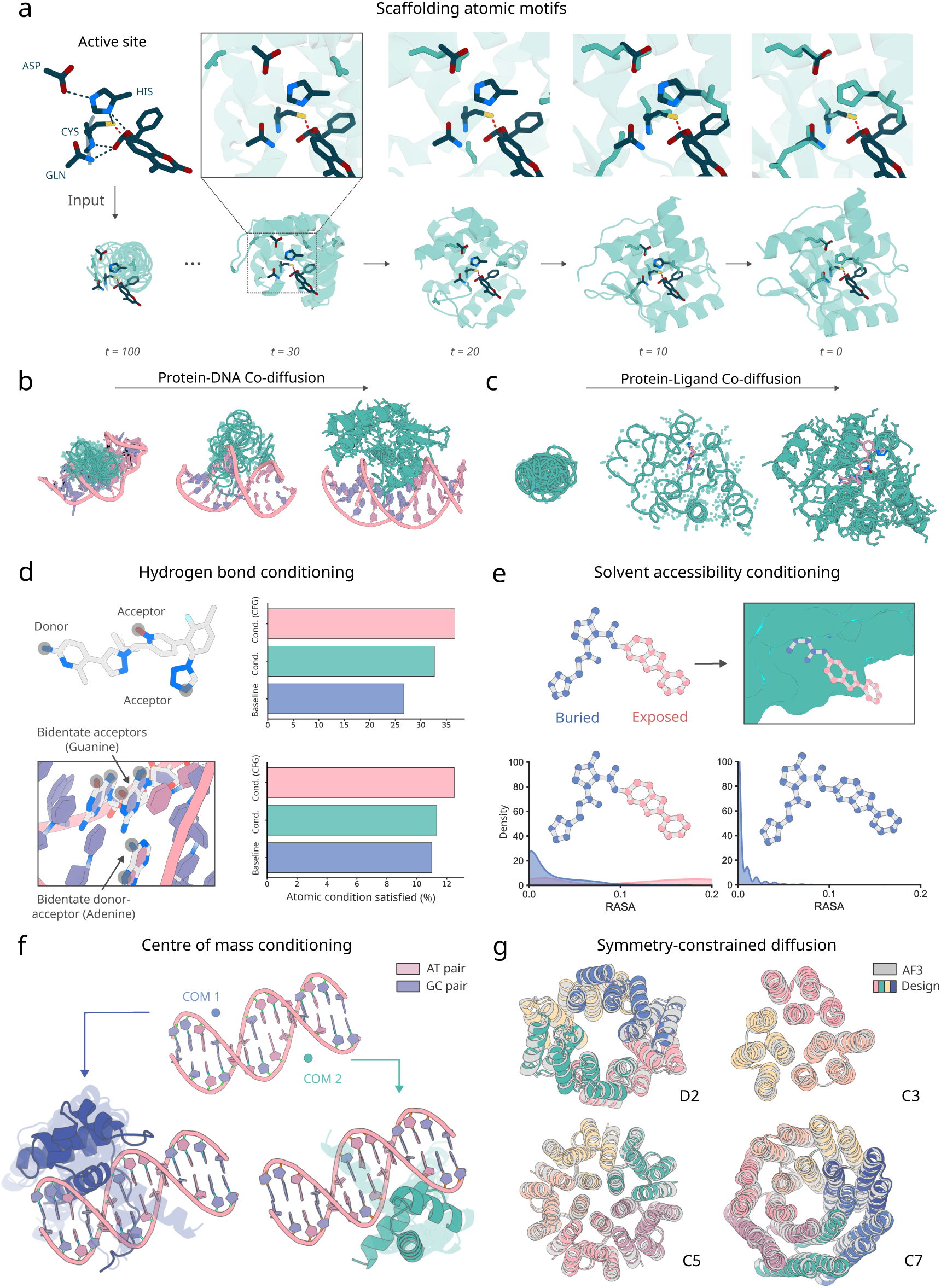
Global and atom-level conditioning with RFdiffusion3. **a**, Diffusion of enzyme active sites. Sidechains are diffused for all residues of the backbone, the optimal position in sequence to scaffold the atomic motif is thereby found during generation, allowing active sites to be scaffolded from simple active site constraints. (top) Zoom in of a trajectory showing the network deciding where in the sequence the side chain residues belong. (bottom) Visualization of predicted denoised structures across different time scales. **b,** Depiction of a diffusion trajectory where both the protein and the DNA are co-generated. In this case, the DNA sequence is given and only the conformation is sampled. **c,** Depiction of a diffusion trajectory where both the protein and the small molecule coordinates are co-generated. In this case, the ligand identity is given and only the conformation is sampled. **d,** Atom-level hydrogen bond conditioning. Both acceptor and donor atoms can be specified as atom level constraints for ligands (top) and nucleic acids (bottom). Applying classifier-free guidance improves adherence to the conditions in both cases. **e,** Atom-level conditioning of solvent accessibility by relative solvent accessible surface area (RASA). (top) example of atomic specification of accessible surface area input specification and output structure (bottom) distributions of the generated structures with different input specifications (over 400 designs from the small molecule benchmark). **f,** The generated centre-of-mass can be guided by the initialized noise cloud. (top) DNA input with two different protein centres of mass. (bottom) generated structures superimposed (with transparency) cluster around the input centre of mass in each case. **g,** Symmetric scaffolds can be specified at inference time by symmetrizing outputs of the diffusion modules. Shown are D2, C3, C5, and C7 all with AF3 C*α* RMSDs 0.832 Å, 0.450 Å, 0.614 Å, and 0.539 Å respectively.

**Figure 3:**
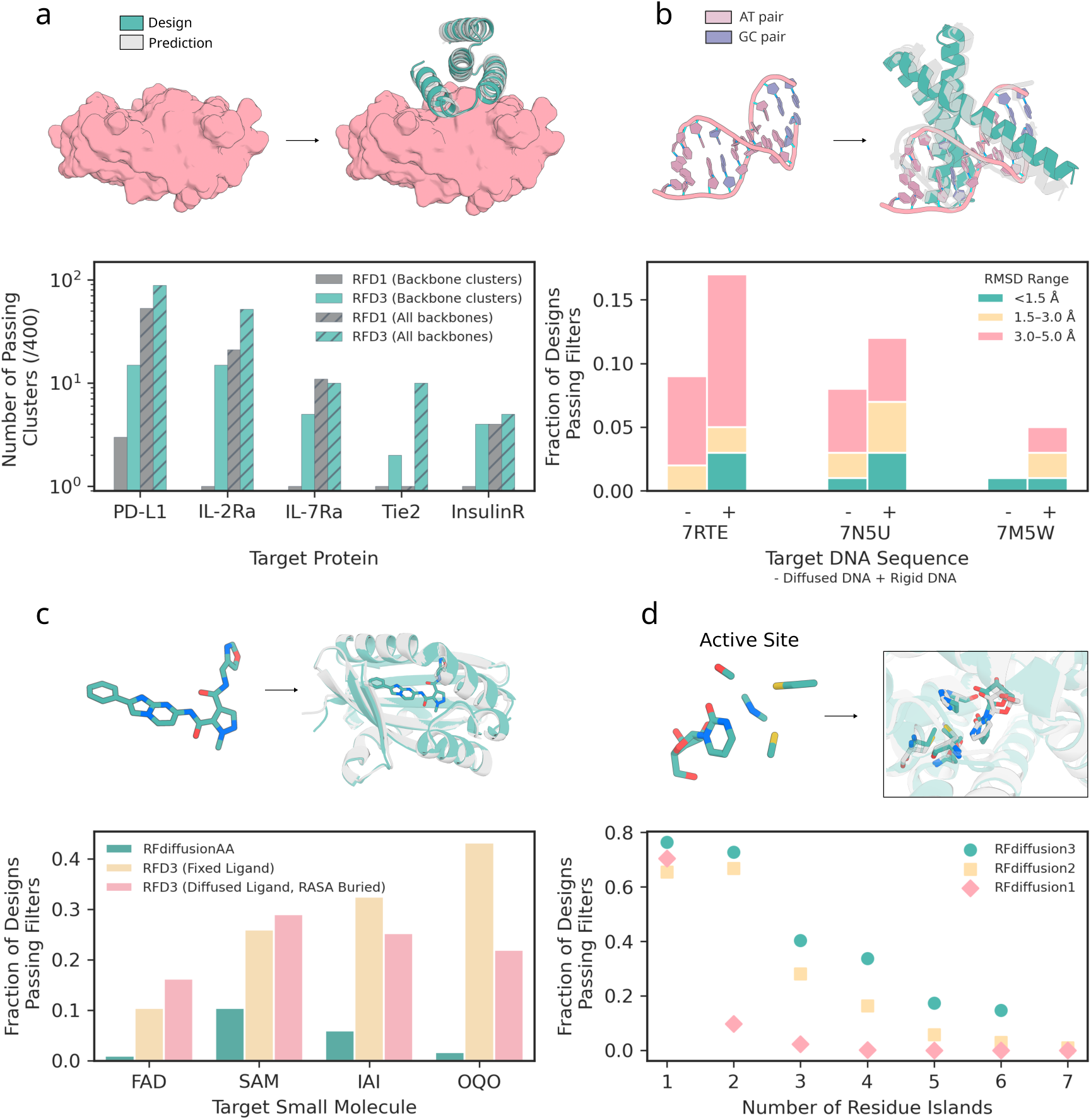
Performance of RFdiffusion3 on *in silico* benchmarks. All panels: Designs shown in teal, predictions shown in gray (prediction method specified in each figure caption). **a,** Generation of protein binding proteins. RFdiffusion3 was compared to RFdiffusion1 (noise scale 0) across five targets (PD-L1, IL-2Ra, IL-7Ra, Tie2, InsulinR). (top) example generation of protein binding-protein to Tie2 (PDB ID: 2GY5; target structure cropped for visualization) with an AF3 prediction overlayed. (bottom) designs that meet AF3 success (min PAE < 1.5, pTM binder > 0.8 and RMSD < 2.5 Å after generating 4 sequences for each backbone with ProteinMPNN) criteria outlined for protein-protein interfaces. Solid bars indicate number of successful clusters (TM score threshold 0.6) and hatched bars represent the total number of successful backbones, both out of 400 total backbones. RFD3 outperforms RFD1 in 4/5 cases without clustering and all cases when clustering, since RFD3 is able to find consistently more diverse solutions to each binding problem. **b,** Generation of DNA-binding proteins. (top) depiction of DNA inputs (PDB ID: 7RTE) and a generated structure with an AF3 prediction overlaid. (bottom) measure of AF3 success (DNA-aligned RMSD) across three DNA targets (PDB ID: 7RTE, 7N5U, 7M5W) across different thresholds (teal < 1.5 Å, yellow 1.5-3 Å, pink 3-5 Å). 400 backbones were generated for each DNA target (and 4 sequences fit to each with LigandMPNN) with both fixed DNA conformation from the input PDB (+) or given only the sequences and predicted the DNA shape along with the protein scaffold (−). The minimum RMSD for each backbone was taken as a representative to score each backbone. **c,** Generation of small molecule binding proteins. Four small molecules are benchmarked (two are common in the PDB; FAD, SAM; and two are uncommon; IAI, OQO). (top) Example of an input ligand (IAI) and a generated structure with the AF3 structure overlayed. (bottom) Comparison of RFD3 to previous method RFdiffusionAA on small molecule generation (400 backbones for each condition; backbones for RFdiffusionAA from published work [2]). AF3 success is defined as backbone aligned ligand RMSD < 5 Å, backbone aligned backbone RMSD < 1.5 Å, min chain pair PAE < 1.5 and iPTM > 0.8. Fraction of designs indicates how many of the 400 generated backbones had at least one sequence that passed the criteria described. Results for RFdiffusionAA shown in teal, RFD3 with fixed ligand using a PDB conformation in yellow and RFD3 with diffused ligand and RASA conditioning (described in Figure 2) in pink. **d,** Generation of enzymes. (top) Example scaffolding of a 4-residue active site (M0097; PDB ID: 1CTT). (bottom) Performance across the Atomic Motif Enzyme (AME) benchmark with increasing number of residue islands. Residue islands are defined as the number of contiguous regions in sequence space that are provided as input to scaffold. Passing backbones are defined as those where at least one of 8 LigandMPNN sequences have a Chai-1 predicted motif backbone-aligned motif all-atom RMSD < 1.5 Å.

We trained on all complexes in the Protein Data Bank (PDB) [22, 23] including protein-protein, protein-small molecule, and protein-DNA interface design tasks, as well as scaffolding functional motifs (Table S1) and a set of high quality AlphaFold2 (AF2) [24] distillation structures predicted in Hsu et al. [25]. Each training structure is annotated with features that designers could potentially specify such as hydrogen bonds, binding partners, or functional motifs. Training examples are generated by noising native protein structures to different extents and predicting the positions of the backbone and side chain atoms for a given residue provided some subset of the applicable conditioning information. For all amino acids except for tryptophan, the extra unused atoms are placed on the C*_β_* position of the residue (C*_α_* for glycine). In our formulation, the terminal oxygen of serine and the sulfur of cysteine are different virtual atoms, thus allowing the network to distinguish between those otherwise identical residues. Given the range of tasks for training RFD3, we adopt a hierarchical training procedure to prevent overfitting. We pre-train the network on a mix of AF2 predictions and the structures deposited in the PDB, and then fine-tune on a larger fraction of DNA and PPI. When training across the PDB, we use entries deposited through December 2024. Full details of the architecture and training procedures are provided in the Supplemental Methods.

## 3 *In silico* results

We first considered unconditional protein generation, in which the model produces novel backbones without specification of a binding partner. The generated structures are diverse (41 clusters out of 96 generations between length 100-250 when using a cutoff TM-score of 0.5) and the amino acid sequence composition of the generated proteins (inferred from the final output coordinates) is similar to the ProteinMPNN generated sequence distribution but has a bias towards alanine compared to native sequence distributions (we hypothesize this is because the network sample distribution is biased towards compact globular folds compared to the PDB) (Fig. S1b). We found that the consistency of the sequence with the structure (as assessed using AF3 or RF3) could be improved by redesign using ProteinMPNN [26], which is not surprising given that RFD3 only sees the final backbone structure in the last diffusion step. Therefore, in all the tests below, we used ProteinMPNN (or LigandMPNN [27] for challenges with non protein atoms) to generate sequences after completing the diffusion trajectory.

To evaluate unconditional protein generation, we designed proteins for chain lengths between 100 and 200. We find that 98% of designs have at least one sequence that is predicted by AF3 to fold within 1.5 Å RMSD to the backbone of the design model (out of 8 generated sequences by ProteinMPNN). Since RFD3 utilizes sparse attention and has a lean architecture, the model scales significantly better than previous approaches using diffusion (Fig. 1d), achieving approximately a 10-fold increase in speed over RFdiffusion2 for typical protein length ranges. In the following sections, we analyze the ability of RFD3 to generate proteins for different design challenges.

### 3.1 Design of protein-binding proteins

Deep learning methods have made impressive improvements in protein binder design, enabling identification of binders from screening fewer than 96 proteins [1, 28, 8, 29, 30]. As in previous work, we enable users to specify the desired binding epitope using a “hotspot” feature; while previous methods define “hotspots” at the residue level, RFD3 enables users to specify atom-level “hotspots” for more precise control over desired interactions.

We benchmarked RFD3 against RFD1, a commonly used binder design method. We generated 400 designs against five therapeutically relevant targets with each method: Programmed Death-Ligand 1 (PD-L1), Insulin Receptor (InsulinR), Interleukin-7 Receptor-*α* (IL-7Ra), Angiopoietin-1 Receptor (Tie2), and Interleukin-2 Receptor-*α* (IL-2Ra). After sequence fitting with ProteinMPNN, we evaluate the designs using AF3 prediction confidence cutoffs from [8] (minimum inter-chain pAE ≤ 1.5; binder pTM ≥ 0.8; target-aligned binder C*_α_* RMSD *<* 2.5Å). RFD3 outperforms RFD1 for 4/5 targets when considering all backbones; notably, when clustering the generated backbones (by TM-score 0.6), we find that RFD3 finds significantly more unique solutions to each binding problem (Fig. 3; on average 8.2 vs 1.4 unique successful clusters for RFD3 and RFD1, respectively; see Fig. S2 for more extensive analysis of characteristics of *in silico* success). The diversity is not limited to the monomer structure, as we find that RFD3 also samples more diverse docking poses than RFD1 (Fig. S2f). We expect that RFD3 will complement existing tools for protein binder design, especially for cases with low *in silico* success rates.

### 3.2 Design of protein-nucleic acid interactions

*De novo* design of nucleic acid binding proteins is a grand challenge in protein design. Existing design methods are primarily family-specific (Zinc fingers [31], TALE [32]). Previous efforts to *de novo* design nucleic acid binding proteins involve docking idealized nucleic acid binding motifs into pre-computed scaffold libraries [33]. Design success rates for nucleic acids are often low [33], due to their polar and flexible nature [34, 35]. We hypothesized that just as protein-only generative diffusion methods have enabled generating protein binding proteins, a network that explicitly modeled nucleic acids could generate DNA binding proteins.

A key challenge when designing nucleic acid binding proteins is inferring the bound DNA shape of the target sequence [36, 37, 38]. To address this, we trained the network to jointly predict the protein structure and the DNA conformation, given the sequence we want to bind (Fig. 2b). We tested DNA binder generation on three DNA sequences that were excluded from the training set. For each sequence, we generate 100 structures using RFdiffusion3, assign four sequences to each backbone using LigandMPNN [27], and then predict their structures using AF3. To assess the accuracy of the design, we align the DNA phosphate atoms of the design and the prediction and compute the RMSD of the protein C*_α_* atoms while trimming terminal loops on the N and C terminus (DNA-aligned RMSD) (Fig. 3b). We found that removing the pre-specification of the DNA structure slightly reduces the success of refolding for monomeric generation (Fig. 3b) but samples more diverse DNA conformations (Fig. S3g). For dimeric generation, removing pre-specification of DNA structure improves refolding success in some cases (Fig. S3d). On average, RFD3 achieves a pass rate of 8.67% for monomeric designs and 6.67% for dimeric designs (*<* 5Å DNA-aligned RMSD). When the RFD3 generated interface is fixed during LigandMPNN sequence design, observed pass rates are 6.5% for monomeric designs and 5.5% for dimeric designs (Fig. S3cd).

### 3.3 Designing small molecule binders

Proteins that bind small molecules have broad applicability as sensors, diagnostics and light harvesting particles [39, 40, 41, 42, 43, 44, 45]. Previous diffusion-based methods have enabled the design of proteins capable of binding small molecules but require the input of a rigid ligand conformation for binding. Similar to flexible DNA, we hypothesize that the atomic nature of RFD3 would enable jointly sampling the protein structure and the target ligand geometry.

We benchmarked RFD3 on four diverse molecules studied in [2] using AF3 prediction of the structure of the small molecule-protein complex to measure success (protein backbone RMSD ≤ 1.5 Å & backbone-aligned ligand RMSD ≤ 5 Å & Interface min PAE ≤ 1.5 & ipTM ≥ 0.8). We found that RFD3 significantly outperforms RFdiffusionAA in all four cases when the ligand was kept rigid (Fig. 3c). We then explore the additional ability of RFD3 to jointly generate the small molecule coordinates and find the same trend when specifying a buried RASA label with classifier free guidance. We also find that the RFD3 designs are more diverse, novel compared to the training set, and have lower binding energies by Rosetta ΔΔG calculations. The ability to sample the ligand conformation jointly with the protein structure surpasses previous methods where a rigid conformation was necessary and will simplify design for ligands with several potential conformers.

### 3.4 Enzyme design

Designing *de novo* enzymes requires precisely arranging specific atoms to catalyze a reaction [46, 47, 48, 49, 50]. RFD2 overcame the limitations of having to hard code the sequence position and rotamer of the catalytic residues but relied on a hybrid residue-atom representation that involved co-diffusing both backbone frames and catalytic residue atoms simultaneously [3]. We hypothesized that the fully atomistic representation of RFD3 would significantly reduce the complexity of generating scaffolds around minimal atomic motifs.

To scaffold unindexed atomic motifs with RFD3, we append an extra token for every catalytic residue to the diffused tokens, allowing the network to process the active site constraints in context. As opposed to the diffused tokens which each contain 14 atoms, the atomic motif conditioning tokens only correspond to the atoms that are being fixed in the atom-level transformer. The model is trained to overlap one of the diffused protein side chains with the fixed atoms (Fig. 2a).

We compared RFD3 to RFD2 on the atomic motif enzyme (AME) benchmark, which measures the ability of networks to generate proteins that house specific side chain constraints on 41 active sites from the PDB as measured by Chai, an open-source reproduction of AlphaFold3 [51]. We find that RFD3 outperforms RFD2 on 37 of the 41 cases (90%) (Fig. S5c). In RFD2, we found that cases with more than four residue islands (separate contiguous stretches of residues) were difficult to scaffold. However, RFD3 yields significantly better solutions to cases with over 4 residue islands (15% vs 4% passing designs for RFD3 and RFD2, respectively; *n* = 12 cases). Additionally, RFD3 produces novel and diverse folds (Fig. S5). We further subsetted the AME enzymes into examples with C2 symmetry and scaffolded the symmetric motif across chains using the native symmetric transform. To measure success, we use AlphaFold3 because of its improved accuracy [11]. When initialized with symmetric noise, we found that RFD3 was able to generate scaffolds that house the active site geometry in both subunits in all cases, including a difficult case with seven residue islands (Fig. S6).

## 4 Experiments

We hypothesized that strong performance of RFD3 on *in silico* benchmarks would translate to improved ability of the network to generate functional proteins *in vitro*. We selected two design challenges that involve the atom-level conditioning capabilities of RFD3, DNA-binding and enzyme design, to highlight the capabilities of the network.

To design DNA binding proteins, we follow a two stage protocol. First, we sample designs directly from the model conditioned on an AF3 prediction of a randomly generated target DNA sequence. Second, for well-predicted designs, we fix the motif contacting the DNA and resample the rest of the backbone to further optimize the structures. We obtained synthetic genes encoding 5 designs, measured binding affinity using yeast surface display and flow cytometry, and found one bound with an EC50 of 5.89 ± 2.15 *µ*M (Fig. 4; Fig. S7 for detailed description of the design process).

**Figure 4:**
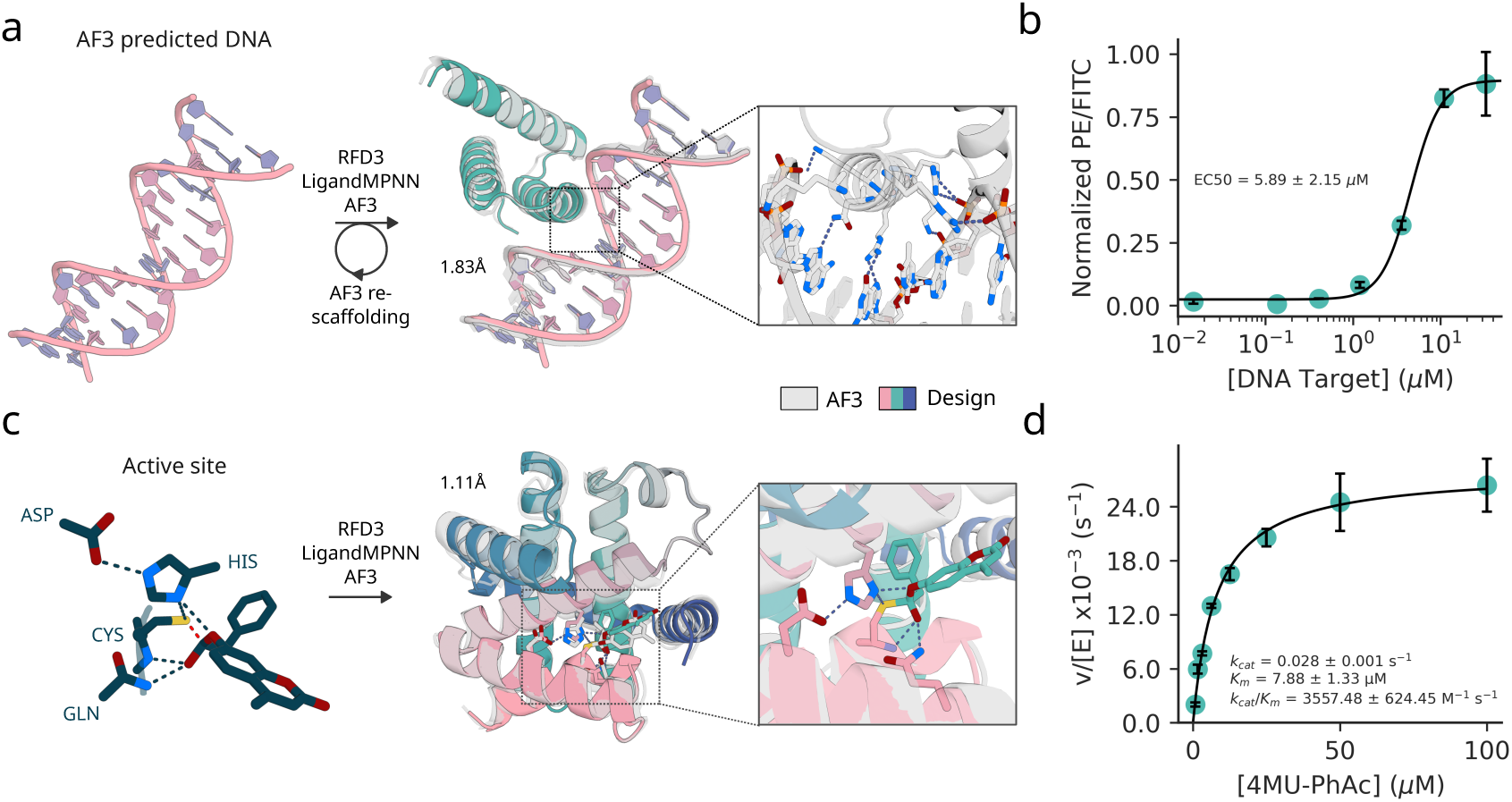
Experimental evaluation of RFdiffusion3. **a,** Schematic representation of DNA binding protein design with RFdiffusion3. Binders were diffused against a fixed DNA structure of the target sequence CGAGAACATAGTCG and LigandMPNN was used to design sequences. Major groove recognition regions of AF3 passing designs were re-scaffolded to increase diversity. Full details available at supplementary section Section 4.2. Structure shown for validated design DBRFD3, polar interaction in the major groove are highlighted (AF3 prediction). **b,** Normalized binding signal (PE/FITC) from flow cytometry data of yeast surface display based titration assay (without avidity) revealed a low micromolar binding affinity of DBRFD3 (1 out 5 designs tested against the target sequence). The average and standard deviation of the EC50 value (computed by fitting a four-parameter logistic regression model) is reported based on three replicates. **c,** (left to right) input active site for the cysteine hydrolase design, design structure overlayed with AF3 model, zoom of active site geometry. **d,** Michaelis-Menten kinetics of designed Cysteine Hydrolase.

For enzyme design, we chose an esterase reaction where a Cys-His-Asp catalytic triad enables hydrolysis of an activated substrate (4-methylumbelliferyl phenyl acetate) as a first step towards the broader class of esterase reactions which have therapeutic and industrial applications[52, 53, 54]. As in RFD2, we defined a minimal motif using the crystal structure of a native cysteine hydrolase (Ulp-1, PDB ID: [55]) comprising the active functional groups of the canonical Cys-His-Asp triad, Gln, the backbone atoms of the residues flanking the cysteine, and the substrate positioned in the first tetrahedral-intermediate geometry. We screened 190 designs and found 35 multi-turnover designs with the most active enzyme exhibiting a *K_cat_/K_m_*= 3557, exceeding previous designs for the same reaction (Fig. S8) [3, 56].

The experimental evaluation shows that the improved atomic conditioning in RFD3 enables improved precision of design for prospective design challenges. In addition to improving the precision of conditioning information, the simultaneous generation of protein backbone and sidechains enables the backbone to better encode residues that make productive atomic contacts with target molecules (small molecules, DNA, etc). To show this, we test whether MPNN designs interactions with target molecules at the same residue positions and whether those residue positions are also conserved for charge with the RFD3 input sidechain and find that MPNN recapitulates many of the designed interactions (Fig. S9a-d). Finally, we find that the designed interaction residue types are often biophysically plausible and that fixing these interactions and only redesigning the remaining residues does not significantly affect the refolding metrics (Section 3.3c-d, Fig. S4f).

## 5 Conclusion

RFD3 outperforms previous methods [1, 2, 3] *in silico* on a wide range of protein-protein binding, protein-DNA binding, protein-small molecule binding, and enzyme design tasks, at a fraction of the compute cost. This improved performance has several origins. First, atom-level design constraints are most naturally implemented in a model where the atoms are the fundamental unit, rather than residues. For RFD2 [3], the specified “tip” atoms were essentially a new data type, as all other components are residues; in contrast, in RFD3 the tip atoms are identical to all other tokens except that their coordinates are fixed (Fig. S5a). Furthermore, in RFD2 general atom-level conditioning on hydrogen bond donor and acceptor status and atomic burial cannot be carried out since only a small subset of atoms are represented explicitly. Second, while our 14 atom per residue approach has been experimented with previously, this was done in the context of architectures more similar to the AF3 pairformer and so were more compute intensive, and no approaches were implemented for providing the conditioning information critical to designing proteins with new functions. The RFD3 network is much faster than both the RFD1 [1] and RFD2 networks, which inherited much of the RoseTTAFold [57, 2] architecture, but at the same time RFD3 is much more expressive as conditions can be placed on every atom in the system.

The control that can now be achieved in the design of complex functions with RFD3 is unprecedented: for example, an enzyme active site can be specified by providing coordinates for a subset of substrate/transition state atoms and the surrounding sidechain atoms; the hydrogen bonding donor/acceptor status of others (often for catalysis what matters is that a protein sidechain donates or accepts a proton from the substrate, but not the identity of that sidechain); the extent of burial of individual sidechain and substrate atoms; and the overall placement of the substrate and active site relative to the protein center of mass. Our extensive *in silico* and experimental tests demonstrate the broad applicability of the method. RFdiffusion3 provides scientists with a general platform for designing binding proteins for essentially any target molecule, catalysts for arbitrary chemical reactions, and complex protein assemblies.

## Acknowledgements

We thank Luki Goldschmidt and the IT team, Kandise VanWormer and the lab management team, for maintaining the computational and wet-lab resources at the Institute for Protein Design. We thank Ian Haydon for developing helpful visualizations of diffusion trajectories and Zari Magness for administrative assistance. We thank Woody Ahern and Arnav Nagle for thoughtful insights during the early development of this project. We thank Bowen Jing for helpful discussions about generative modeling during development and Long Tran for discussions on the best AF3 filters to use for small molecule binding. We thank Magnus Bauer, Julia Bonzanini, Nathan Greenwood, Isaac Sappington, Adam Chazin-Gray, Shayan Sadre, Brian Belmont, Inna Goreshnik, and Linna An for providing feedback from using earlier versions of the network. We thank the Rosetta Commons Hope Woods, Rachel Clune, Rocco Moretti, Sergey Lyskov.

## Author Contributions

Research design: R.K., J.B., D.B.; J.B. led the development of the model architecture; Development and *in silico* evaluation of RFD3 for — DNA binders: R.M., P.K. — Small-molecule binders: Y.L., J.C. — Protein-protein interaction: R.B. — Symmetric oligomers: A.M., H.E. — Enzymes: J.B., J.F.; Development of evaluation pipeline infrastructure: J.B., R.B., J.F.; Built infrastructure that substantially accelerated development: N.C., S.M.; Design and *in vitro* characterization of: — Cysteine Hydrolases: S.S. — DNA Binders: R.M., Y.P., E.S.; Supervision and organization: D.B, R.K., J.B.; Model and code contribution: J.B., N.C., R.B., Y.L., S.M., P.K., J.F., T.T., R.M., H.E., A.M., J.C., F.D., R.K.; Writing — original draft and figures: R.K., J.B., R.B., R.M., J.F., Y.L., J.C., P.K. — review and editing: All authors.

## Supplementary Figures

**Fig. S1:**
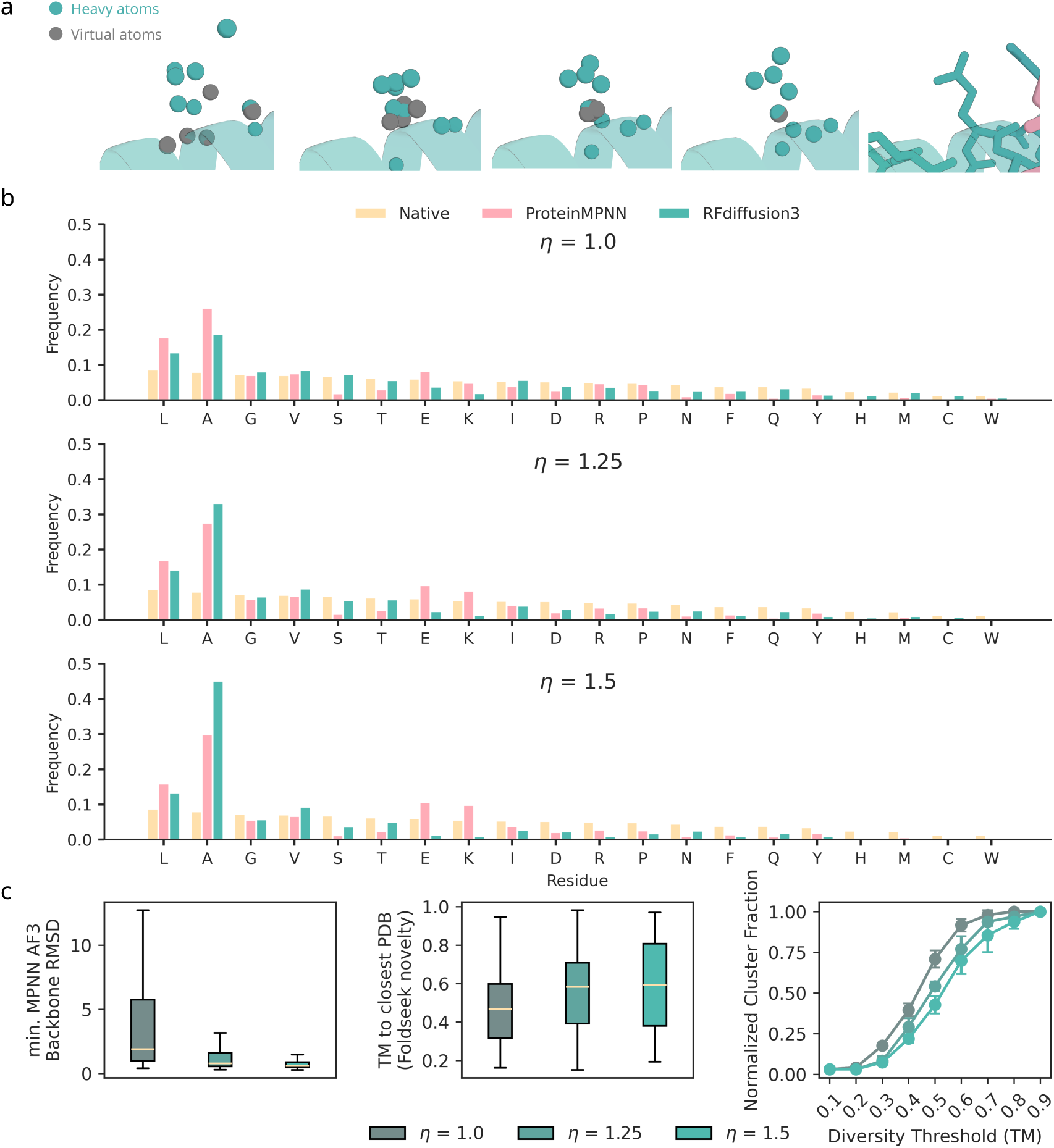
Evaluation of sequence distribution of RFdiffusion3. **a**, Diffusion of sequence via 14-atom representation. Additional virtual atoms are identified after diffusion as those placed on the C*_β_* atom, and the residue identity can be decoded by the atom names of the heavy atoms. **b**, Sequence distributions of RFdiffusion3 for varying step scales (*η*), compared with the native (PDB) distribution, and the ProteinMPNN sequence fitted to the backbone. **c**, (left-to-right) Unconditional metrics of RFdiffusion3: designability of RFD3 with step scale (*η*); Novelty of RFD3 with step scale; Number of clusters under different TM thresholds as measure of diversity, normalized to the total number of clusters at TM= 1. Overall, designability increases and diversity decreases with increasing step scale, forming a useful inference parameter for problems of interest. For this work, we opt to use *η* = 1.5 for benchmarks.

**Fig. S2:**
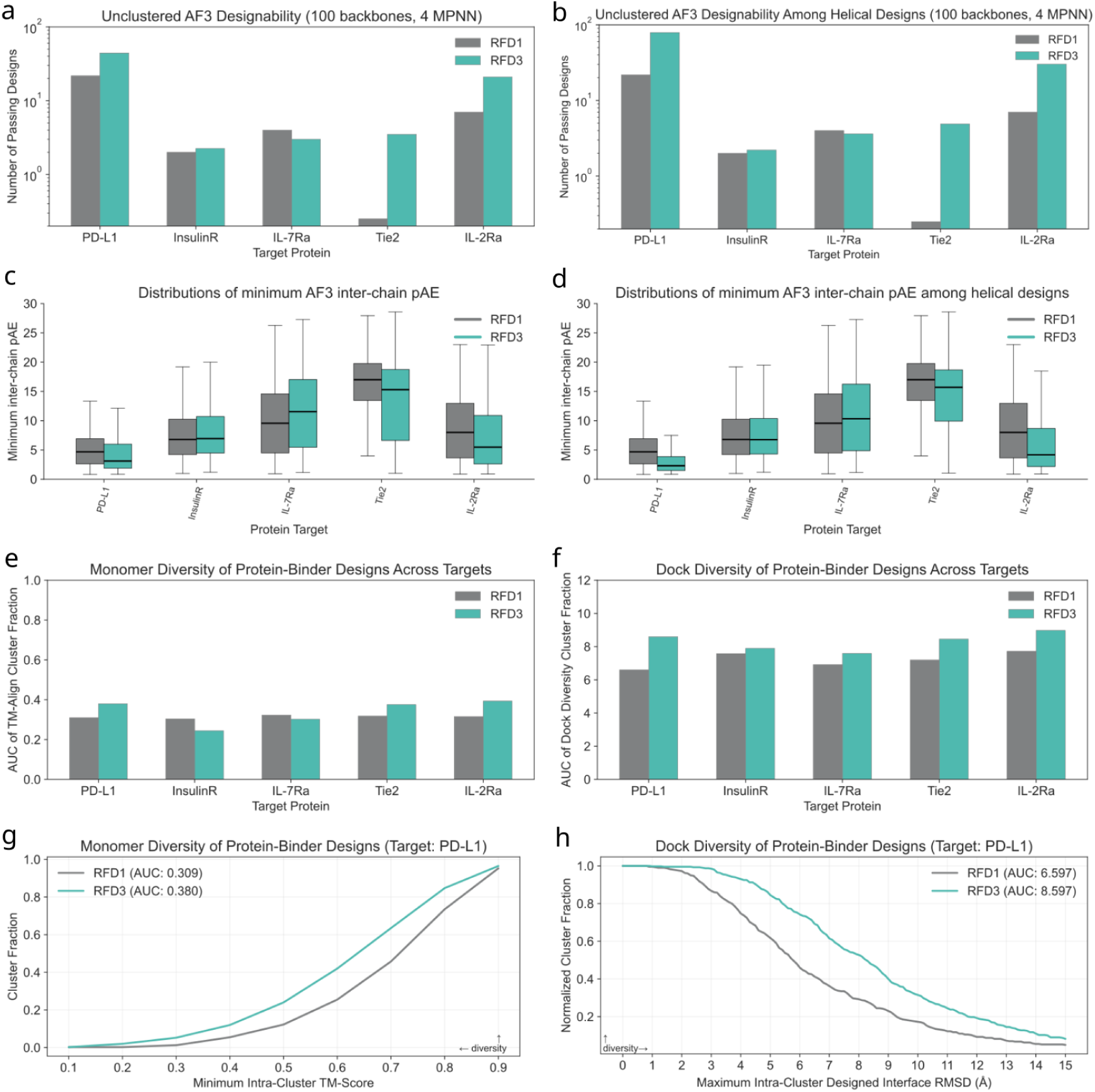
Designability and diversity of PPI Designs with RFD1 and RFD3. 400 backbones were designed for each of the five targets, then ProteinMPNN was used to design 4 sequences per backbone. **a,** Unclustered pass rate among all designs, normalized to 100 backbones (with 4 MPNNs) per target. Note that we use the stricter designability cutoffs introduced in [8] and detailed in the main text. **b,** Unclustered pass rate among primarily helical designs, defined as those with >75% helix content. This included all RFD1 designs and the following per-target counts with RFD3: PD-L1, 142 (36%); InsulinR, 359 (90%); IL-7Ra, 339, (85%); Tie2, 274 (69%); IL-2Ra, 143 (36%). As before, this plot is normalized to 100 backbones (with 4 MPNNs) per target. **c-d,** Distributions of the minimum AF3 inter-chain pAE among all designs and helical designs, respectively. Taken together, these results show that RFD3 significantly outperforms RFD1 when using the full set of designs, and that the advantage grows even further when subsetting to RFD1- style mostly-helical designs. Despite the higher pass rates among helical bundles, we believe it advantageous to have a model that is not limited to this fold type. **e,** AUC values for monomer diversity. For each target, 400 designed backbones were agglomeratively clustered at different TM-score thresholds using the “complete linkage” criterion, meaning that any given pair of elements in a cluster must have a TM-score above the threshold value. The metric used to compute the AUC is “cluster fraction”, defined as the number of clusters divided by the number of examples, calculated at intervals of 0.1 between 0 and 1, exclusive. RFD3 shows higher monomer diversity than RFD1 across 3 of 5 targets. **f,** AUC values for the cluster fraction determined by a new metric, the “dock diversity”. We define the dock diversity as the target-aligned backbone N, C*α*, C RMSD between the 15 residues in the design that are closest to the target by C*α*-C*α* distance. The cluster thresholds are evenly spaced at intervals of 0.1 Å between 0 and 15, inclusive. As before, 400 designed backbones are clustered agglomeratively using the “complete linkage” criterion. RFD3 shows higher dock diversity than RFD1 on all targets. **g-h,** AUC curves for the 400 designs for PD-L1, displaying monomer diversity (**g**) and dock diversity (**h**).

**Fig. S3:**
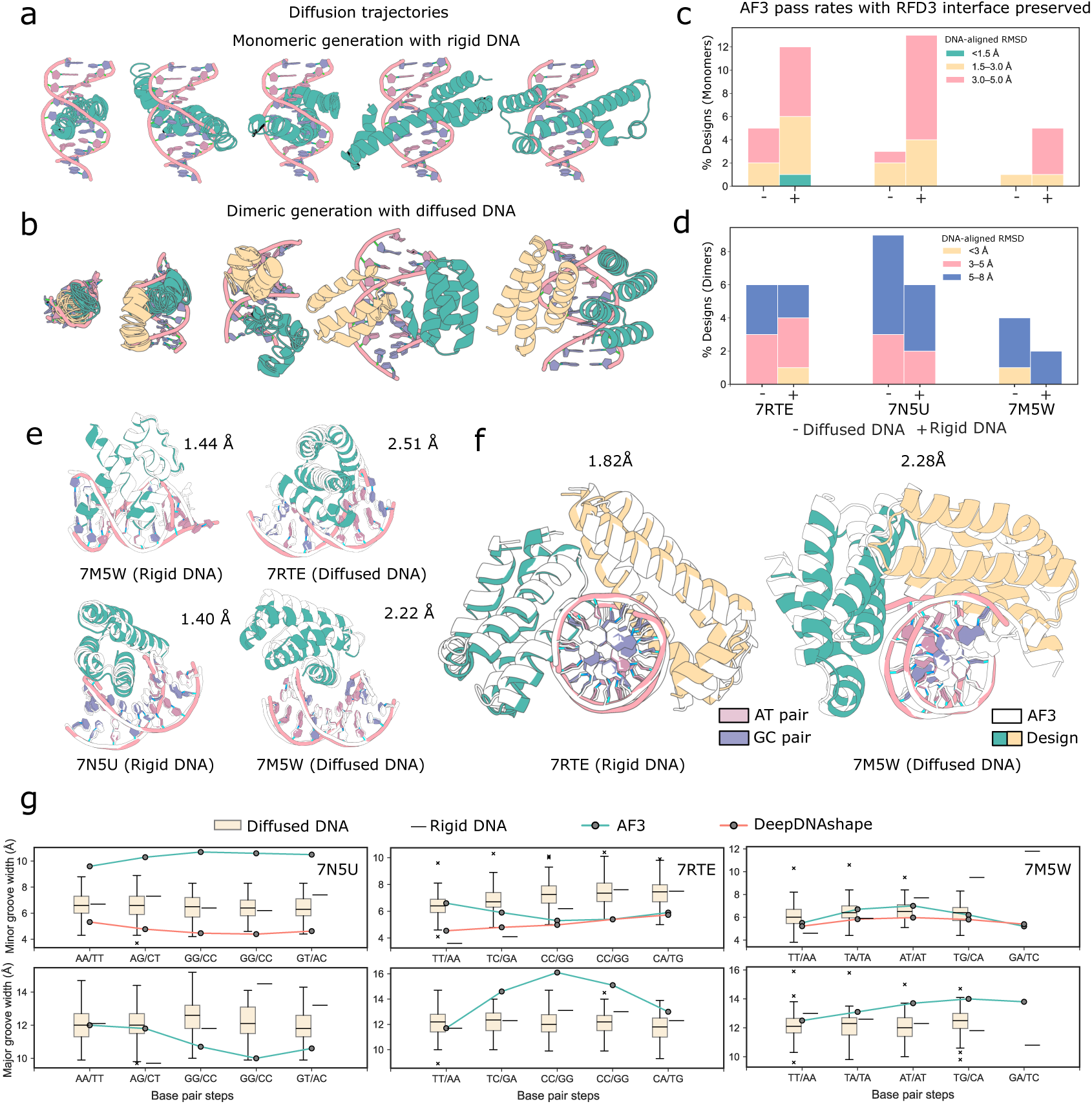
Design of DNA interactions with RFdiffusion3. **a,** Diffusion trajectory of monomeric de novo DNA binder generation using RFD3 against rigid DNA structure (from PDB ID: 7RTE). **b,** Diffusion trajectory of dimeric de novo DNA binder generation using RFD3 against diffused DNA (i.e. RFD3 generates DNA structure; DNA sequence from PDB ID: 7N5U). **c,** Percentage of agreement with AF3 prediction of RFD3 interface preserved (residues within 2-3.5 Å of DNA atoms)-LigandMPNN designs (monomeric) across three different targets, at different DNA-aligned RMSD ranges (*<*1.5 Å, 1.5-3 Å, 3-5 Å) and against both diffused and rigid DNA targets. **d,** Percentage of agreement with AF3 prediction of RFD3 interface preserved (residues within 2-3.5 Å of DNA atoms)-LigandMPNN sequences (dimeric) across three different targets, at different DNA-aligned RMSD ranges (*<*3 Å, 3-5 Å, 5-8 Å) and against both diffused and rigid DNA targets. **e,** Illustration of agreement with AF3 predictions for selected monomeric designs. **f,** Illustration of agreement with AF3 predictions for selected dimeric designs. **g,** RFD3 generated diverse protein-bound DNA conformations in the context of corresponding PDB (rigid) conformations and oracle predicted (AF3 and DeepDNAshape [58]) free DNA conformations. Major (bottom row) and Minor (top row) groove widths were computed using 3DNA-Analyze [59] for RFD3 generated (diffused and rigid target) and AF3 predicted structure of three different DNA target sequences. DeepDNAshape predicted minor groove width is also shown (averaged over consecutive bases to achieve base-pairstep values consistent with 3DNA-Analyze. Major groove width prediction was unavailable) Note: for 7M5W, output of 3DNA-analyze only contained values for 4 base-pair steps.

**Fig. S4:**
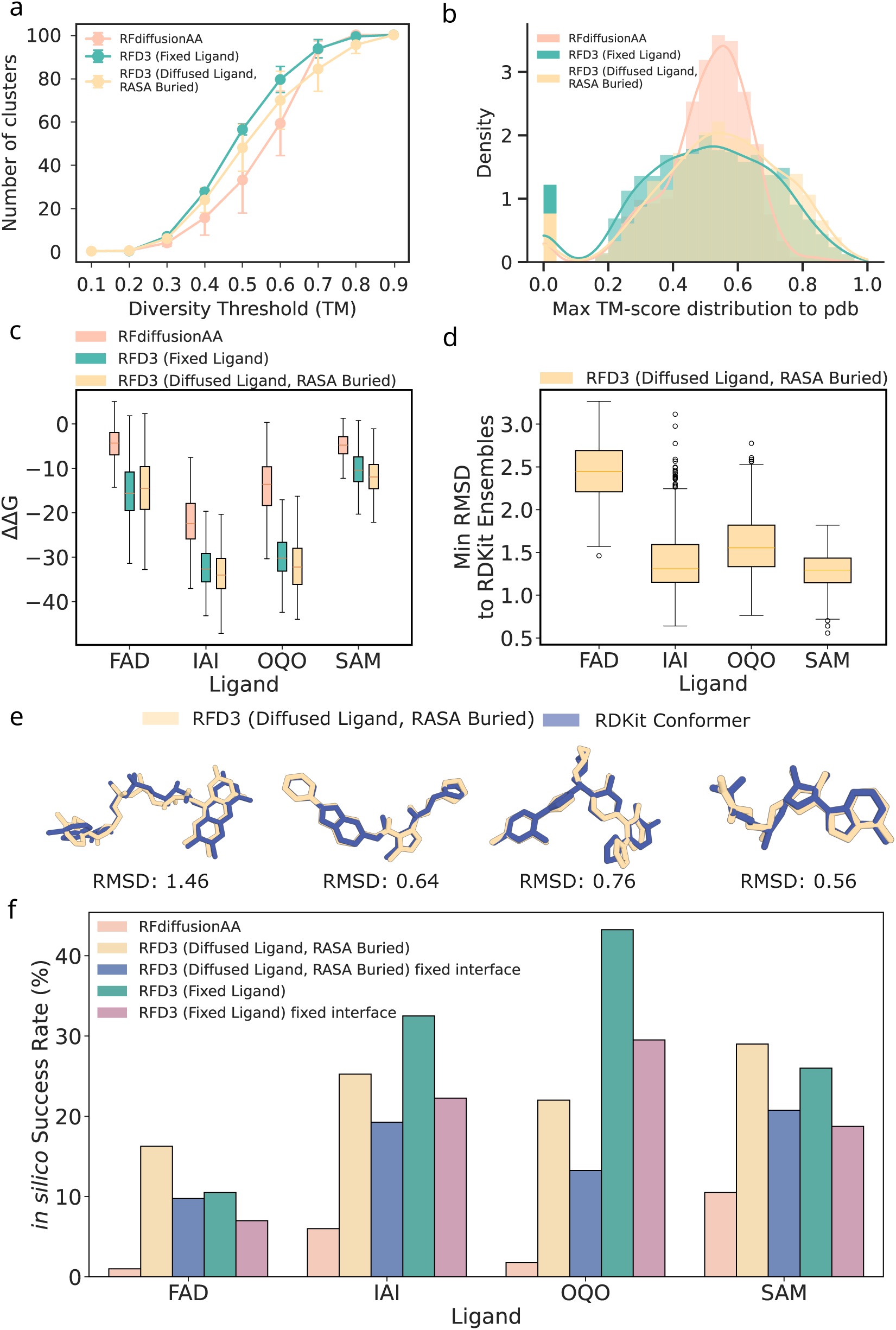
Design of small-molecule binding proteins with RFdiffusion3. **a,** Diversity of generated small molecule binders. Pairwise structural similarity among the designs is assessed using all-by-all TM-align. To quantify diversity, we performed agglomerative clustering with TM-score thresholds ranging from 0 to 1 and reported the number of clusters at each threshold. For comparison, the same analysis was applied to 400 designs generated by RFdiffusion All-Atom for each target. Results across four ligands were averaged, with error bars representing variability across targets. Overall, RFdiffusion3 generated more diverse structures than RFdiffusion All-Atom. **b,** For the same set of 400 RFdiffusion3 designs described in *a*, we used FoldSeek to identify the closest structural neighbor in the PDB for each binder and recorded its TM-score. We then evaluated novelty by counting the number of designs whose maximum TM-score (i.e., similarity to the closest PDB structure) falls below thresholds ranging from 0 to 1. This quantifies how distinct the generated binders are relative to known protein structures. The analysis was performed for each method, aggregated across all four ligands. **c,** We evaluated the binding energies of the designs described in *a* using Rosetta, applying the DDG_norepack_metric to assess binding quality. This metric estimates the change in binding free energy (ΔΔ*G*) between the bound and unbound states of a complex, without side-chain repacking during the calculation. Lower values indicate stronger binding. Outliers have been excluded to improve the clarity of the visualization. **d,** RMSD distribution between RFdiffusion3-generated diffused ligands and RDKit conformers. For each ligand target, 50 conformers were generated using RDKit. For each design under the diffused-ligand setting, we computed the RMSD between the generated ligand and all conformers and reported the minimum RMSD. **e,** Representative structures of RFdiffusion-generated diffused ligands showing the smallest RMSD to RDKit-generated conformers. **f,** Comparison of *in silico* designability across different settings. In addition to the standard benchmark, where LigandMPNN is used to redesign the entire binder sequence, we also assessed designability under a fixed-interface setting, in which interface residue identities were preserved and only the remaining regions were redesigned with LigandMPNN. The ligand–protein interface is defined as protein residues within 2–3.5 Å of any ligand atom. All evaluations use 400 backbones and 8 sequences per design, and the shared inference parameters of step scale *η* = 1.5, noise level *γ*_0_ = 0.6, and 200 denoising steps (used throughout this work unless otherwise specified.) Finally, diffused-ligand binders were generated with the RASA condition set to buried and a CFG scale of 2 to guide the creation of ligand binding pockets.

**Fig. S5:**
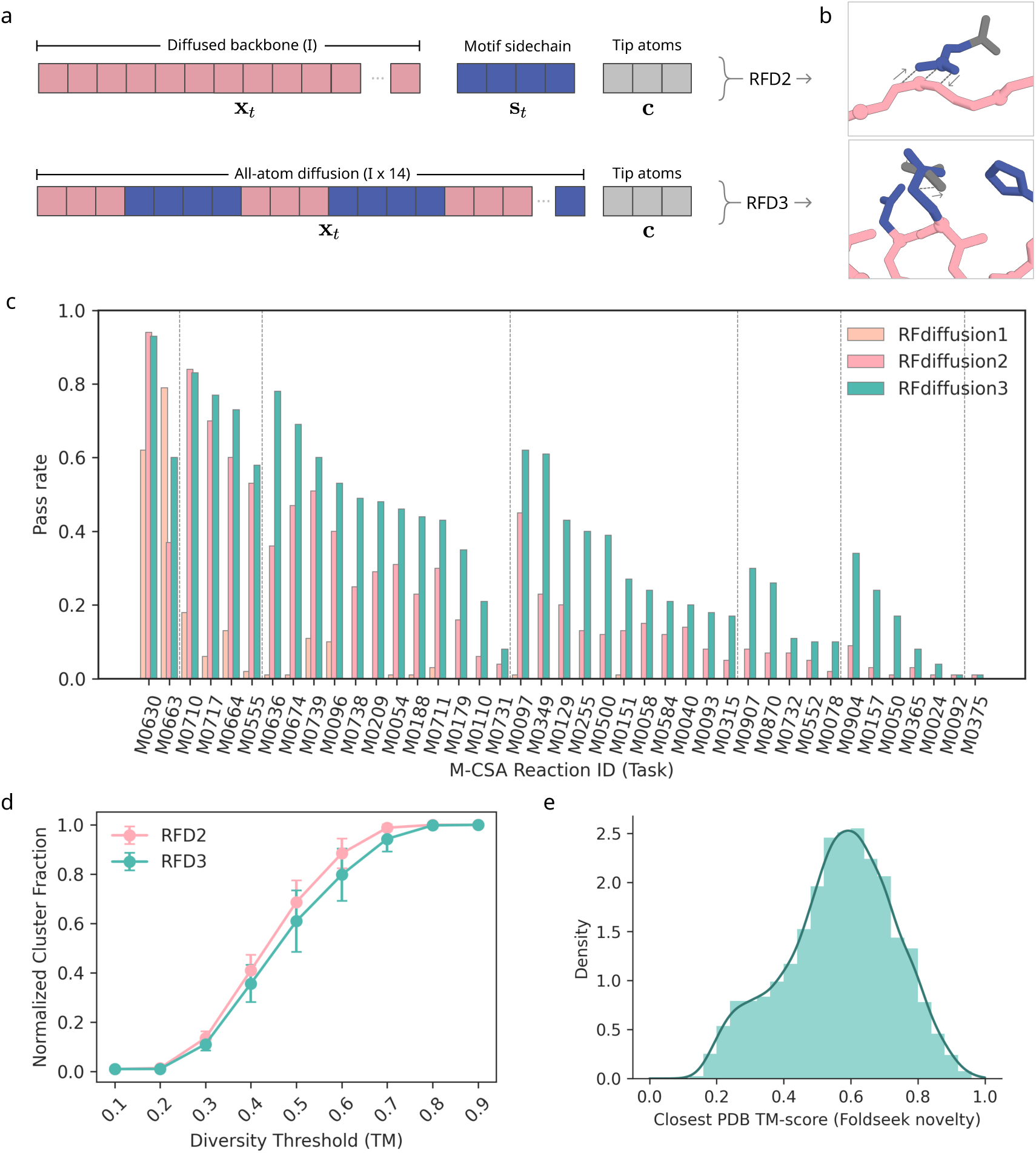
Design of enzymes with RFdiffusion3. **a,** Atomic Motif Enzyme Benchmark of RFdiffusion variants. For benchmark details see section Section 3.5. **b,** Tokenization of unindexed atoms with RFdiffusion2 involves explicit tokenization of the motif sidechains belonging to the (fixed condition) tip atoms. In contrast, RFdiffusion3 natively diffuses sidechains, making explicit tokenization of motif sidechains unnecessary. **c,** Diversity analysis of RFD3 against RFD2. Diversity analysis includes full generated ensemble. Error bars indicate the variance across the 41 different AME tasks at a particular diversity threshold. **d,** Novelty analysis of outputs on the AME benchmark. RFD3 produces a significant number of novel folds (signified by high density between TM 0.2 and 0.4.)

**Fig. S6:**
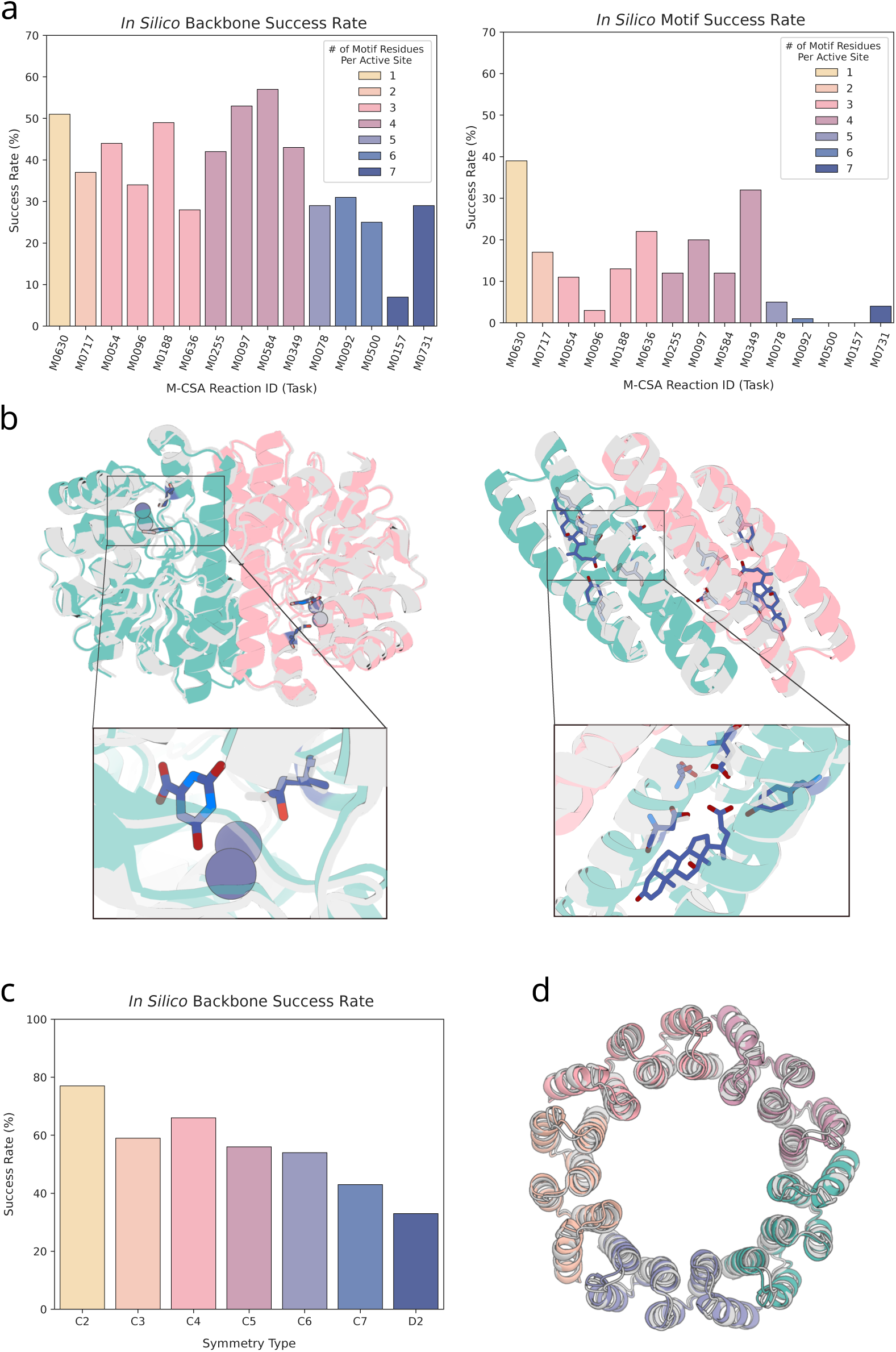
Design of symmetric enzymes with RFdiffusion3. **a,** *In silico* success rates for backbones on a subset of the AME benchmark. (left) Success rate for backbones of complexes. (right) Success rate of active site residues in an asymmetric unit (ASU) on top of the complex backbone success. **b,** Example outputs from **a**. (left) M0630 C2 enzyme with complex backbone C*α* RMSD of 0.84 Åand motif all-atom RMSD of 0.46 Å(AF3). (right) M0349 C2 enzyme with complex backbone C*α* RMSD of 1.08 Å and ASU motif all-atom RMSD of 0.66 Å. **c,** *In silico* success rate for backbones on symmetry-constrained diffusion without motifs. **d,** Example output for C5 symmetry with C*α* RMSD of 0.91 Å.

**Fig. S7:**
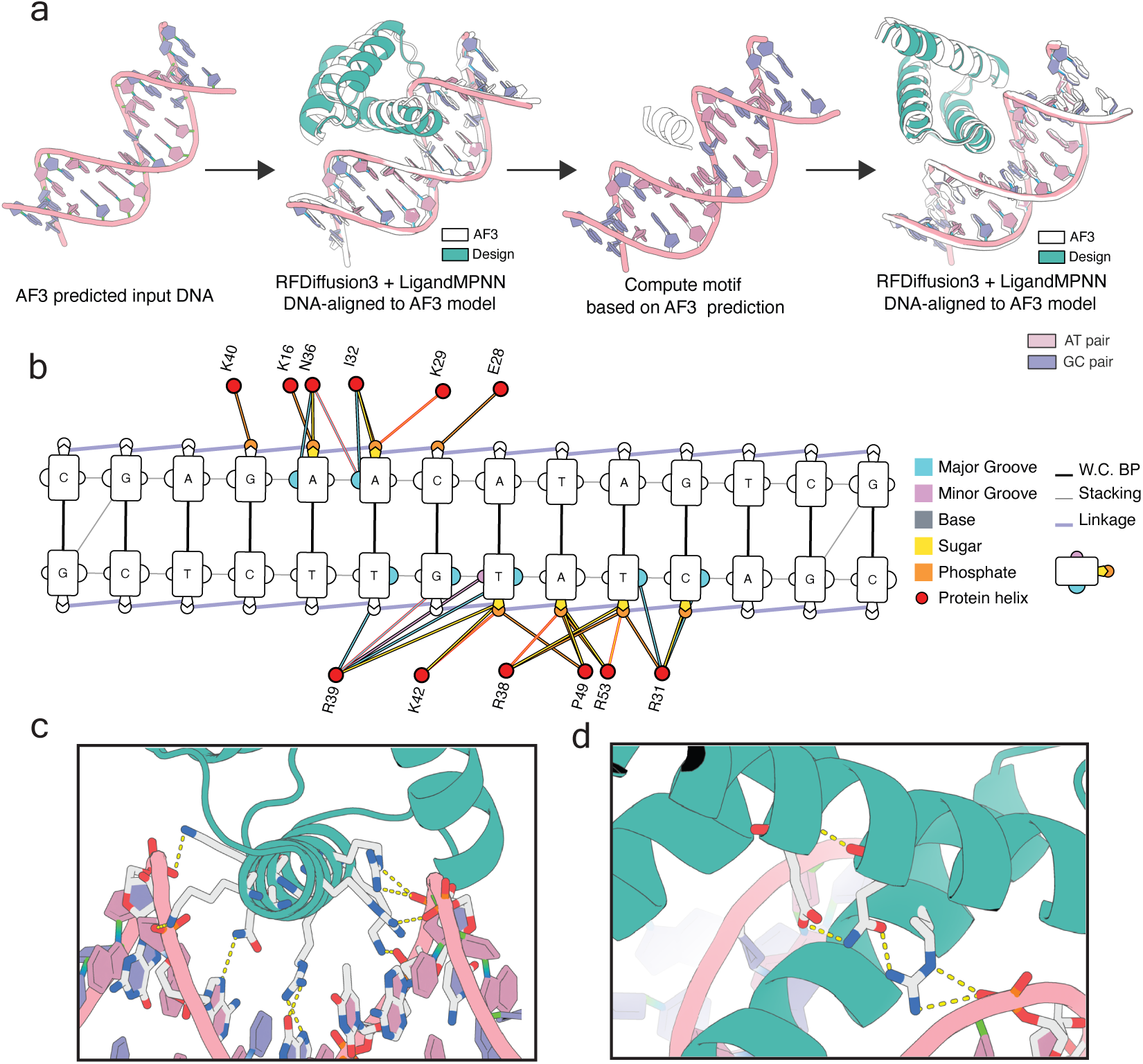
Design of DNA binders with RFdiffusion3. **a,** Schematic representation of the DNA-binder design pipeline used to design DBRFD3. Full details in Section 4.2. **b,** Protein-DNA contact map of DBRFD3 generated using DNAproDB [60]. Interaction lines indicated with bold red borders denote hydrogen bonding. **c,** DBRFD3 recognizes DNA through multiple major groove and backbone interactions with ASN36 (N36) and ARG39 (R39) performing major groove hydrogen bond interactions, to recognize adenine and guanine bases, respectively. **d,** Hierarchical supportive hydrogen bond interactions observed in DBRFD3 (usually uncommon in *de novo* designs). The positively charged residue ARG38 interacts with the negatively charged phosphate moiety. The neutral residue GLN56 supports ARG38. GLN56 is, in turn, supported by negatively charged GLU60.

**Fig. S8:**
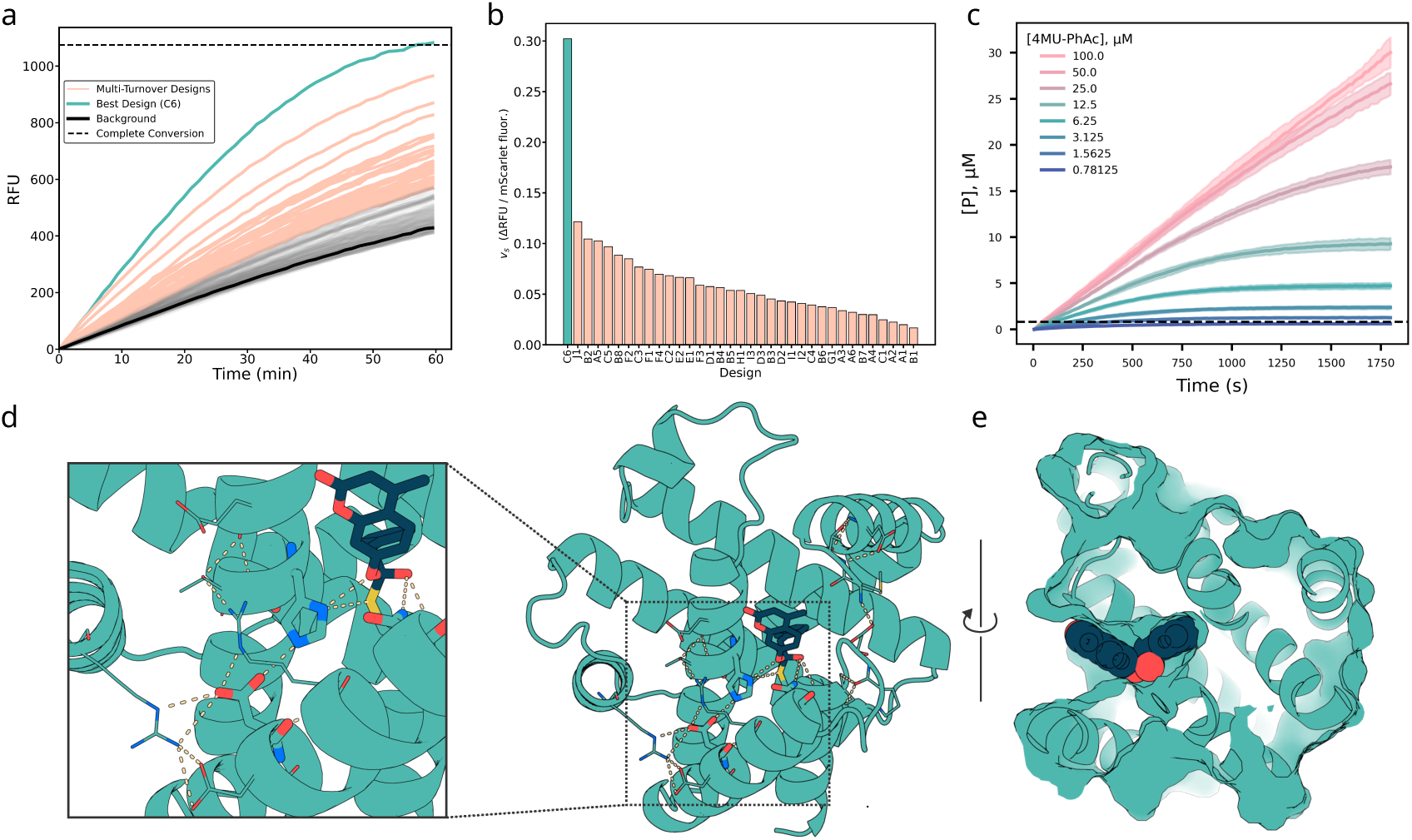
Experimental characterization and structural analysis of *de novo* cysteine hydrolase designs. **a,** IVTT screen of 190 designs using 4MU–PhAc substrate. Representative fluorescence traces (RFU) are shown for background controls (black), multi-turnover designs exceeding the 10% conversion threshold (cyan), and the best design, C6 (orange). The dashed line indicates full conversion. **b,** Normalized steady-state rates (background-subtracted and scaled by mScarlet fluorescence) for the 35 multi-turnover designs, grouped by scaffold family and indexed numerically. C6 is highlighted in orange. **c,** Michaelis–Menten kinetics of purified C6 measured in E. coli expression. Product formation is shown across eight substrate concentrations (mean ± s.d., n = 3). Dashed line shows enzyme concentration in the reaction well (0.8 *µ*M) **d,** Active site of C6 (AF3 prediction). (left) zoomed-in view showing pre-organized hydrogen-bonding network at the active site: the Asp general base locked in place by dual hydrogen bonds from two arginines, themselves supported by Asp and Thr a nearby helices. (right) Global view of scaffold with catalytic residues and substrate in blue, and the hydrogen-bonding networks highlighted. **e,** AF3 prediction of C6 in complex with the modeled tetrahedral intermediate (TI1), showing shape-complementary pocket geometry and correct orientation of the 4MU leaving group toward solvent.

**Fig. S9:**
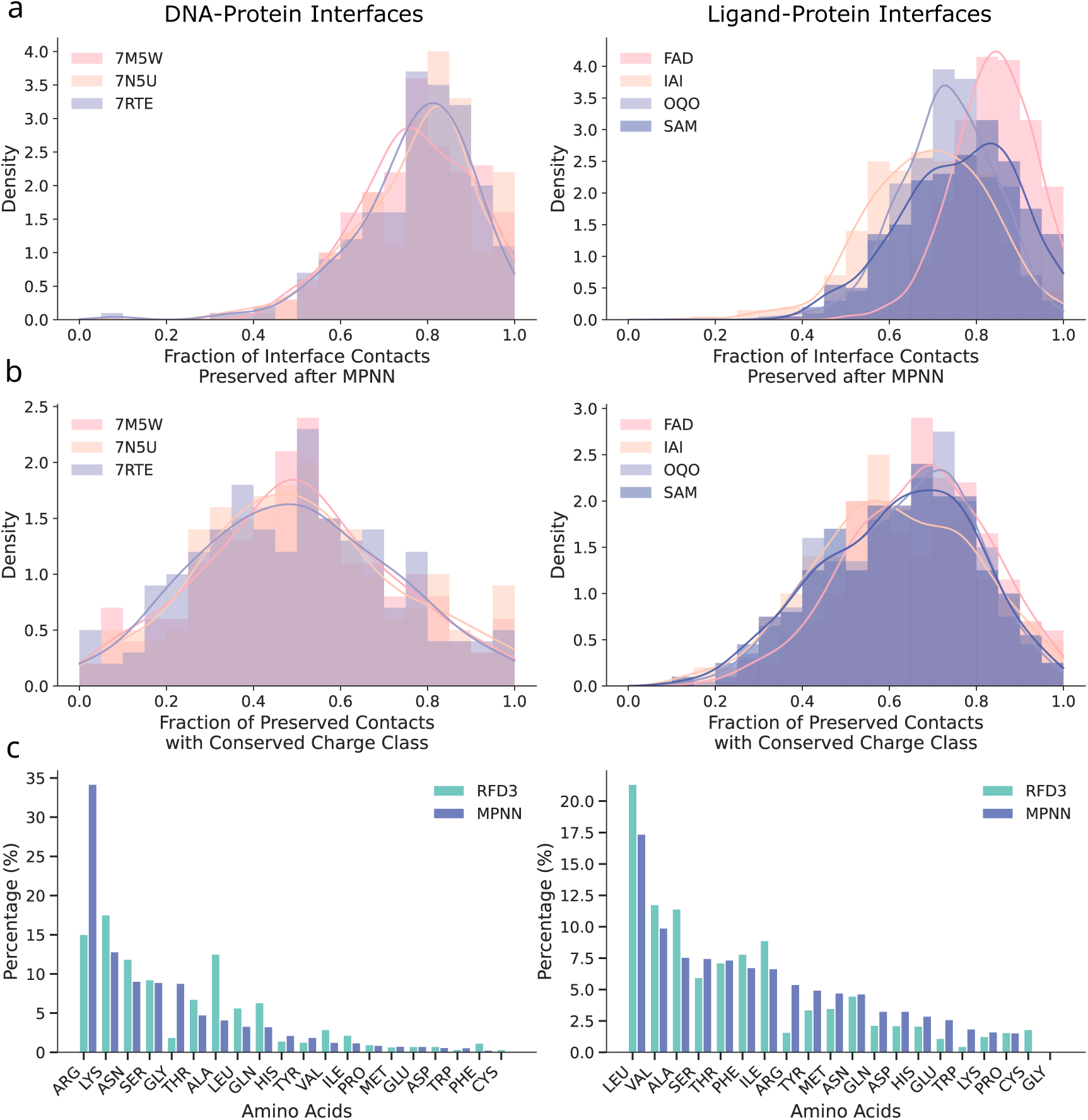
Analysis of interface residue composition between RFdiffusion3 generated sequences, LigandMPNN sequences and native sequences. **a**, Fraction of interface residue contacts preserved after MPNN. For each design case, we generated 400 structures using RFdiffusion3. For DNA targets, ProteinMPNN was used to design 4 sequences per structure, and for ligand targets, LigandMPNN was used to design 8 sequences per structure. Interface residues were identified in both the RFdiffusion3 designs and the corresponding MPNN-designed structures using a 2–3.5 Ådistance threshold. We then computed, for each design, the fraction of interface contacts that were preserved before and after MPNN sequence design, and reported the average value across all the MPNN-packed structures for each design. **b**, Fraction of preserved interface-contact residues that also retain their charge class (positive, negative, or neutral). Using the same setting as in *a* we measured, among all preserved interface contacts, the fraction where the contacting residues share the same charge class before and after MPNN. **c**, Distribution of amino acids at the interface. Interface residues were defined as protein residues within 2–3.5 Åof any DNA or ligand atom.

## Footnotes

1 1github.com/RosettaCommons/atomworks

