## Supplementary Information for "*De novo* Design of All-atom Biomolecular Interactions with RFdiffusion3"

for

**1 RFdiffusion3 Training Regimen** **47**

### Supplementary Methods

#### Notation

| | |
|---|---|
| **General terminology** | |
| Token | Standard amino acid and nucleotide residues are represented as single tokens. Modified residues, all ligands, and unindexed conditions are tokenized per-atom. |
| Indexed | Tokens which *have* global residue index information revealed |
| Unindexed | Tokens which *do not* have residue index revealed |
| **General notation** | |
| $I$ | Number of tokens for sample |
| $L$ | Number of atoms for sample |
| $A$ | Maximum number of atoms per token for sample |
| $D$ | Diffusion batch size |
| **Common Tensors and Shapes** | |
| $\mathbf{q}_l \in \mathbb{R}^{D\times L\times c_{\text{atom}}}$ | Processed atom-level features |
| $\mathbf{c}_l \in \mathbb{R}^{L\times c_{\text{atom}}}$ | Encoded atom-level conditioning features |
| $\mathbf{p}_{lm} \in \mathbb{R}^{L\times L\times c_z}$ | Encoded atom-pairwise features |
| $\mathbf{a}_i \in \mathbb{R}^{D\times I\times c_{\text{token}}}$ | Processed token-level features |
| $\mathbf{s}_i \in \mathbb{R}^{I\times c_s}$ | Encoded token-level conditioning features |
| $\mathbf{z}_{ij} \in \mathbb{R}^{I\times I\times c_z}$ | Encoded token-pairwise features |
| $\hat{\mathbf{x}}_0 \in \mathbb{R}^{D\times L\times 3}$ | Predicted denoised structure |
| $\hat{\mathbf{x}}_l^{\text{self}} \in \mathbb{R}^{D\times L\times 3}$ | Self conditioned structure |
| $\mathbf{x}_l^{\text{noisy}} \in \mathbb{R}^{D\times L\times 3}$ | Noisy input coordinates |

### 1 RFdiffusion3 Training Regimen

#### 1.1 Diffusion training and inference

Similar to Ref. 11, we adopt the EDM [60] diffusion framework. Notably, in contrast to AF3, we do not align predictions from the model after each denoising step:

$$L_{\text{total}} = \alpha(\sigma)\frac{1}{3\cdot|w_l|}\sum_l w_l||\hat{\mathbf{x}}_0^l - \mathbf{x}_0^{\text{gt},l}||_2 + \alpha_{\text{lddt}}\, L_{\text{smooth\_lddt}}, \tag{1}$$

where $L_{\text{smooth\_lddt}}$ is equivalent to the AF3 implementation of smoothed lDDT, $\alpha_{\text{lddt}} = 0.25$, data variance $\sigma_{\text{data}} = 16$, and $\alpha(\sigma) = \frac{1+(\sigma/\sigma_{\text{data}})^2}{(\sigma\cdot\sigma_{\text{data}})^2}$. During training, we sample the noise level according to $\sigma = \sigma_{\text{data}} \cdot \exp(-1.2 + 1.5 \cdot \mathcal{N}(0, I))$.

**Loss weights.** The per-atom loss weights $w_l$ are zero for unresolved atoms. To encourage prioritisation of the unindexed regions, we increase the loss on these tokens at low noise. We do this using by substituting the following value for $w_l$ in Eq. (1): $w^l_{\text{unindexed}} = w^l \cdot \alpha(\gamma\sigma)/\alpha(\sigma)$, applied to tokens which correspond to the unindexed conditioning in the ground truth. We set $\gamma = 0.75$. Additionally, we up-weight the loss on ligand tokens by a factor of 10. Virtual and non-virtual atoms are treated equally in the loss.

**Inference.** During inference, we sample noise levels according to: $\sigma = \sigma_{\text{data}} \cdot (s_{\text{max}}^{1/\rho} + t \cdot (s_{\text{min}}^{1/\rho} - s_{\text{max}}^{1/\rho}))^\rho$, where $s_{\text{min}} = 4 \cdot 10^{-4}$, $s_{\text{max}} = 160$, $\rho = 7$ where $t$ decreases uniformly from 1 to 0 over 200 timesteps.

**Optimizer and training parameters.** For the optimizer, we use AdamW with a learning rate of 1.8e−3, with $\beta_0 = 0.9$ and $\beta_1 = 0.95$. We use EMA with a decay rate of 0.999, and all inference is performed using the shadow weights. We use a linear learning rate schedule, with 1000 warmup steps from 0. We add 10% dropout to all outputs of the processed attention blocks, excluding the atom-level SparseTransformer applied during the encoding phase (before recycling). We train RFdiffusion3 on 16 NVIDIA H200 GPUs, which took approximately 7 days.

#### 1.2 Tokenization pipeline

##### 1.2.1 Atom14 and all-atom tokenization

Each residue is padded to 14 atoms, where additional virtual atoms are placed on the CB. Thus, each residue token contains 14 atoms. Nucleic acids are provided with corresponding numbers of atoms for each token.

For residues, virtual atoms are distinguished through the embedding of the atom names which serves as a form of positional encoding to the network. The atom name encoding is generated based on the scheme shown in Fig. S10. Since cysteine and serine are structurally near-identical, we disambiguate their ordering based on the virtual atom encoding. To do so, we permute the input atom name to the network during training, which is equivalent to permuting the ground-truth, provided the network does not have a means of determining the original canonical order. The network does not use the conventional AF3 sequence-local attention, since the windowed attention would have the scope to leak the number of atomic neighbours, and thereby the permuted sequence.

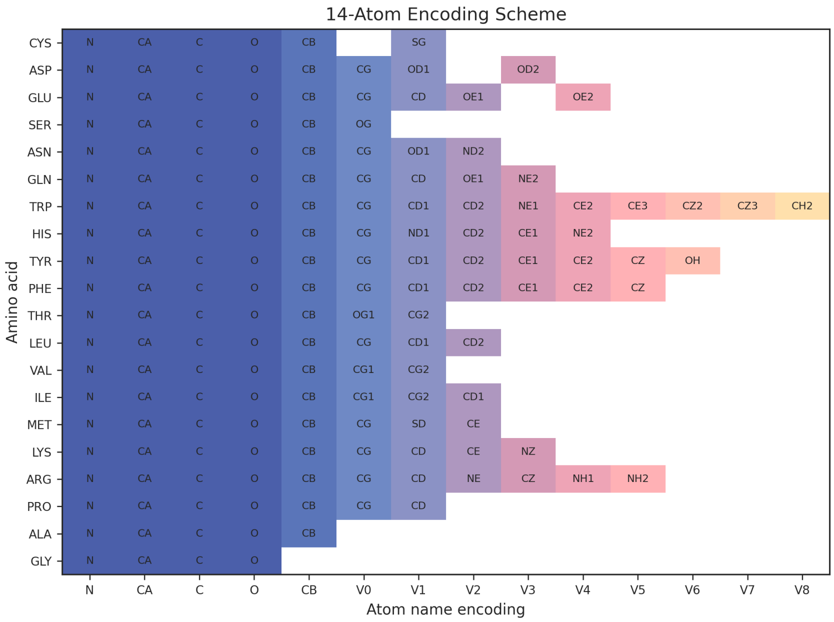

**Fig. S10:** Virtual atom name encoding scheme applied within the RFD3 dataloading pipeline to disambiguate atom orderings.

##### 1.2.2 Indexed tokenization

In the context of protein design, *indexed* tokens refer to those tokens whose sequence positions are known relative to the entire structure. Indexed tokens by default have both their locations within the sequence fixed as well as their 3D coordinates. For indexed tokens, users may specify a range of combinations of fixed sequence, fixed coordinates, or specific fixed atoms only. For example, when scaffolding backbone motifs, the user may specify whether they wish to relax the sequence input provided to the model (and whether to pad the residue with virtual atoms).

##### 1.2.3 Unindexed tokenization

*Unindexed* tokens, introduced in [3], are defined as coordinate-conditioned motif residues that have an unspecified position within the output sequence. Unindexed tokens allow for less user pre-specification and as a result yield greater overall diversity of sampling [3]. Such constraints are often useful when solving geometrically difficult motifs, such as within the design of enzymes (Fig. S5).

The all-atom formulation of RFdiffusion3 allows convenient specification of unindexed residues. In contrast to RFD2, the diffused region need not specify explicit additional atoms belonging to the constrained atom (Fig. S5a). Thus, diffusion proceeds via the movement of atoms from a single track of the network (Fig. S5b), and tip atoms are concatenated as additional tokens. Importantly, unindexed residues need not be atomized, and tip-atom functional groups in RFdiffusion3 are processed as a single token comprising multiple atoms. We find that this design helps scale the use of unindexed scaffolding across tasks such as larger contiguous backbone motifs.

To control the information provided to RFD3 about the global offset of a particular unindexed residue, we provide a special *UNK* bin for the relative positional encoding (specified in algorithm 3.)

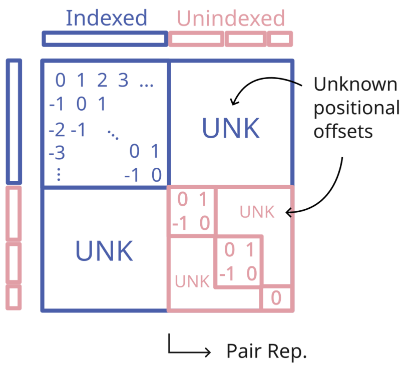

**Fig. S11:** By removing known indices, RFD3's positional encoding enables unindexed tokenization. Regions are considered *indexed* if they contain information defining their position within the protein sequence. *Unindexed* tokens may have relative positional offsets between one another (partial unindexing) and do not have known global offsets (blue off-diagonals). The unindexed tokens are processed *in-context*, i.e., they are concatenated to the models' input and removed after diffusion is complete. By default, RFD3 *inserts* the unindexed coordinates after diffusion is complete via a greedy matching algorithm.

Unindexed tokens are removed from the outputs after diffusion completes. As shown in Fig. S5b, RFD3 diffuses sidechains to overlap with the provided tip atom constraints. During training, we embed the atom names of the unindexed constraints, allowing the network to generate compatible atoms at the corresponding positions (Fig. S10). While the model will approximately respect the specified tip atom constraints, it is often desirable to re-insert the provided tip atom coordinates after denoising. This process requires matching each tip atom constraint with its corresponding diffused atom, identified by finding the nearest diffused atom with the same name.

##### 1.2.4 Sequence design

RFdiffusion3 adopts a relatively straightforward in-context conditioning approach for motifs. For all fixed atoms, we remove the time embedding. Following the EDM formulation of diffusion, at $t = 0$, the model's prediction is guaranteed to exactly recapitulate the input.

#### 1.3 Atom-level conditions

##### 1.3.1 Hydrogen bond conditioning

During training, hydrogen bonds between motif and non-motif regions are identified post-cropping using HBPLUS [61], with a hydrogen–acceptor distance cutoff of 3 Å, donor–acceptor distance cutoff of 3.5 Å, and all other parameters set to default. Motifs may include protein segments, small molecules, tip atoms, or DNA elements.

Donor and acceptor atoms are encoded as atom-level features using two binary scalar channels (`active_donor, active_acceptor`). Motif atoms identified as donors or acceptors are annotated with a value of 1 in the corresponding channel. Non-motif atoms are assigned 0, as they are not available to the model during inference.

To introduce variability and prevent overfitting, we apply *random masking* to the hydrogen bond annotations. Each input has a probability 0.2 of being evaluated for hydrogen bonds; otherwise, all annotations are set to 0. If computed, the donor and acceptor labels are further randomly subsampled with probability 0.5, such that the model is not always exposed to the complete set of hydrogen bond annotations.

##### 1.3.2 Designability conditioning

We include two token-level conditioning features that indirectly control the designability of model outputs: `is_non_loopy` and `ref_plddt`. `is_non_loopy` is a token-level feature, shown with probability 0.99, indicating if the global loop content of the diffused region is $< 30\%$. `ref_plddt` is active for distillation datasets with probability 0.9 and indicates whether the global pLDDT is $> 80$.

##### 1.3.3 Relative accessible surface area (RASA) conditioning

During training, RASA conditioning is applied with a 60% probability when ligands are present. When RASA conditioning is active, the Solvent Accessible Surface Area (SASA) is computed for each atom in the crop using the Shrake-Rupley algorithm [62] implemented in Biotite [63]. The Relative Solvent Accessible Surface Area (RASA) is then calculated by normalizing each atom's SASA value against its theoretical maximum SASA value (computed for an extended conformation), resulting in values ranging from 0 to 1:

$$\text{RASA}_i = \frac{\text{SASA}_i}{\text{SASA}_{i,\max}} \tag{2}$$

where $\text{SASA}_{i,\max} = 4\pi(r_i + r_p)^2$ represents the maximum possible SASA for atom $i$ with van der Waals radius $r_i$ and probe radius $r_p = 1.4$ Å. These RASA values are subsequently discretized into four bins representing distinct solvent accessibility states: "buried" (RASA $\leq 0.06$), "partially-buried" (RASA $\leq 0.2$), "exposed" (RASA $\geq 0.2$), and "unassigned". When RASA conditioning is applied, all ligand atoms are annotated with their corresponding RASA values, motif atoms are annotated with 20 % probability, and remaining atoms retain their unassigned status.

##### 1.3.4 Atom-level PPI hotspots

For PPI examples, the primary form of conditioning is through the provision of atom-level hotspots. In RFD3, hotspots are defined as any atoms on the target that are within 4.5 Åof any atom on the binder, using the same binder/target designations as in the custom PPI crop (Section 1.6). For 75% of such interfaces, a randomly-selected subset of the hotspots in the ground-truth is provided to the model as a binary atom-level feature. When hotspots are provided, the size of the provided subset is sampled uniformly on the interval $[0, \lfloor(n_{gt} * 0.2)\rfloor]$ where $n_{gt}$ is the number of hotspots in the ground-truth.

#### 1.4 Reference Conformer Generation

Inspired by [11], we provide RDKit-generated reference conformers for small molecules (but not standard polymer residues) to the model as an atom-level feature. Following AF-3 methodology, we apply random rigid transformations to each reference conformer to prevent the model from memorizing specific conformations and to improve generalization to different molecular orientations.

We take a number of steps to ensure that these conformers are of high quality, as we find naively parsing structures from the Chemical Component Dictionary (CCD) often results in implausible conformations [14].

Namely, we:

- Add explicit hydrogen atoms to all structures, as force field calculations require proper hydrogen placement for accurate conformer generation
- Re-classify single bonds between metals and coordinating atoms as dative bonds, as we find that the CCD default approach of specifying them as single bonds leads to poor conformer quality
- Apply RDKit's standard molecule sanitization procedures

- *(In the event of further sanitization errors)* Apply ChEMBL's [64] SMIRKS transformations to standardize problematic functional groups (nitro groups, diazonium nitrogens, sulfoxides)
- *(In the event of further sanitization errors)* Apply normalization procedure recommended by RDKit authors
- *(In the event of further sanitization errors)* Compute formal charges based on valence constraints, calculating the difference between electrons in bonds and expected valence electrons
- Generate conformers using RDKit's ETKDGv3 algorithm with UFF force field optimization
- If ETKDGv3 fails (due to excessive constraints or rotatable bonds), fall back to UFF minimization starting from random coordinates
- If all generation attempts fail, use the idealized coordinates from the CCD entry

We also find that conformer generation represents a significant pipeline bottleneck for structures with many rotable bonds, decreasing training speed. To mitigate this challenge, we pre-compute 30 reference conformers for all small molecules in the PDB and sample uniformly from those pre-computed conformers during training.

#### 1.5 Motif centering and input coordinates

Across batches, we apply a random augmentation to the input coordinates:

$$\tilde{\mathbf{x}}^{\mathrm{GT},l} = R \cdot \mathbf{x}^{\mathrm{GT},l} + \sigma_{\mathrm{perturb}}\, z, \tag{3}$$

where $z \sim \mathcal{N}(0, I)$ and $R \in SO(3)$.

The input to RFD3 is centered based on the centre-of-mass of all *diffused* tokens (excluding all fixed tokens). To avoid the model learning to assign special significance to the point $(0, 0, 0)$, we perturb the centre-of-mass in training by a global offset of magnitude $\sigma_{\mathrm{perturb}}$. Further, we augment the noise using a timestep-dependent noise scale, as described by Eq. (4):

$$\hat{\epsilon} = \epsilon + z \cdot \sigma_{\mathrm{perturb\_com}} \cdot \left(\frac{t}{t_{\max}}\right)^{\nu}, \tag{4}$$

where $z \sim \mathcal{N}(0, I) \in \mathbb{R}^3$, $\nu = 3$ and $t_{\max} = 38$. For protein-protein interaction cases, we upweight $\sigma_{\mathrm{perturb}}$ and $\sigma_{\mathrm{perturb\_com}}$ by 2.0x and 1.5x, respectively.

#### 1.6 Cropping

For most training examples, we use one of two standard crops: a Contiguous Crop, which selects group(s) of contiguous residues across chains until the token budget $n_{crop}$ is reached, and a Spatial Crop, which selects the $n_{crop}$ nearest residues to the crop center. To better control memory requirements (since tokens can have variable numbers of atoms), we also introduce a maximum number of atoms in a given crop $n_{atoms}$; otherwise, the details of these two crops are identical to those described in 11. In this work, we set $n_{\mathrm{crop}} = 384$ and $n_{\mathrm{atoms}} = 5000$.

In addition, with probability 0.75, we utilize a custom crop for PPI examples. This custom crop keeps the entirety of one randomly-selected chain (designated as the "binder"), then uses a spatial crop to fill out the remainder of the crop budget. Note that this approach is not used if the binder length exceeds a given threshold `max_binder_length`, in which case the Spatial Crop is used as a fallback. In this work, we set `max_binder_length=256`.

Further, we utilize a custom crop for protein-DNA interfaces, termed the DNA Contact Crop. For this crop, we first sample a protein atom and a DNA base atom that are within 10 Åof each other. Between 15 and 40 protein residues are selected contiguously on either side of the residue containing the protein atom, and between 3 and 10 DNA bases are selected contiguously on either side of the base containing the DNA atom, as well as on the reverse complement strand (if present). This crop produces training examples analogous to DNA binder generation tasks, where protein monomers achieve sequence-specific binding through targeted base interactions with relatively short DNA segments.

The cropping frequencies for each dataset are summarized in Table S3.

#### 1.7 Conditioning sampling

RFdiffusion3 is trained on a unified sampling scheme which selects the tokens and atoms to constrain as the *motif*. During training, each condition (as shown in Table S1) is checked for whether it can be applied to the current example.

| **Parameter** | **tipatom** | **island** | **unconditional** | **sequence_design** |
|---|---|---|---|---|
| `frequency` | 5.0 | 2.0 | 0.0 | 0.1 |
| `island_len_min` | 1 | 1 | 0 | inf |
| `island_len_max` | 1 | 15 | 0 | – |
| `n_islands_min` | 2 | 2 | 0 | – |
| `n_islands_max` | 15 | 5 | 0 | – |
| `p_diffuse_motif_sidechains` | 0.0 | 0.8 | – | 1.0 |
| `p_diffuse_subgraph_atoms` | 1.0 | 0.0 | – | – |
| `residue_p_seed_furthest_from_o` | 0.8 | – | – | – |
| `residue_n_bond_expectation` | 3.0 | – | – | – |
| `residue_p_fix_all` | 0.05 | – | – | – |
| `hetatom_n_bond_expectation` | 8 | – | – | – |
| `hetatom_p_fix_all` | 0.5 | – | – | – |
| `p_fix_motif_sequence` | 0.7 | 0.2 | 0.0 | 0.0 |
| `p_fix_motif_coordinates` | 1.0 | 1.0 | 0.0 | 1.0 |
| `p_unindex_motif_tokens` | 0.95 | 0.0 | 0.0 | 0.0 |

**Table S1:** Probability configuration for different training conditions. Dashed (–) entries indicate unset or unused fields.

#### 1.8 Training Conditions

##### 1.8.1 General Island Class

In this training condition, protein motif tokens are sampled randomly within several "islands", defined as groups of contiguous residues. We sample the number of islands uniformly between 2

| Conditioning type | Frequency |
|---|---|
| `unconditional` | 2.0 |
| `island` | 2.0 |
| `sequence_design` | 0.1/0.5* |
| `tipatom` | 5.0 |
| `ppi` | 0.0/5.0* |
| **Feature** | **Probability** |
| `calculate_hbonds` | 0.20 |
| `calculate_rasa` | 0.60 |
| `keep_protein_motif_rasa` | 0.20 |
| `hbond_subsample` | 0.50 |
| *Unindexed training augmentations* | |
| `unindex_leak_global_index` | 0.20 |
| `unindex_insert_random_break` | 0.33 |
| `unindex_remove_random_break` | 0.33 |
| *Designability conditioning* | |
| `featurize_plddt` | 0.90 |
| `add_global_is_non_loopy_feature` | 0.99 |
| *Protein-protein interaction (PPI)* | |
| `add_ppi_hotspots` | 0.75 |

**Table S2:** Probabilities and frequencies used for meta-conditioning features and training augmentations.
*Different values for training stage1/stage2, respectively

and 5 with 1 to 25 tokens per island, also sampled uniformly. This same strategy also enables unconditional training by setting all of these parameters to 0. Once motif tokens are chosen, further conditioning is assigned based on the probabilities described in Table S1.

#### 1.9 Tip atom sampling

To sample tip atoms in training, we first apply the island mask described above except using only islands of length 1, sampling 2 to 9 islands. We then sample subgraphs of the selected tokens based on their bond graphs as parsed by AtomWorks[14]. The graph is converted to a NetworkX graph and a random seed atom is chosen (with an optional bias towards atoms further from the backbone oxygen). From the seed atom, we sample a connected subgraph by traversing $n \sim \text{Geom}(p)$ bonds within the sidechain structure. The corresponding atoms are used as the tip atoms provided to the model.

#### 1.10 Datasets

Here we report the various datasets used to train RFD3.

**PDB.** The PDB dataset includes structures deposited to the RCSB PDB prior to December 2024. We use the same PDB datasets used to train RF3 [14] (albeit with a different date cutoff); that is, one dataset of chain-like entities (`pn_units`) and one dataset of pairs of interacting chains (`interfaces`). We apply the weighted sampling scheme from [11] to sample from these two datasets, which translates to sampling from the interfaces dataframe $\sim$ 80% of the time and the pn_units dataframe $\sim 20\%$ of the time.

**AFDB Distillation and Interdomain Distillation.** The Alphafold DB Distillation set and interdomain distillation set consist of previously published AlphaFold2 predicted monomers [24] and domain-domain interactions collated from the AlphaFoldDB [65].

**DNA Distillation.** The DNA acid complex distillation set, originally developed for RF3, [14] consists of high-confidence predicted structures of protein-DNA complexes derived from binding pair sequences within experimental data.

**DNA Interfaces.** The protein-DNA interface set is a subset of the PDB interfaces dataset containing all interfaces between a protein and one or more DNA chains. We separate this dataset to facilitate upsampling of these interfaces during the second stage of training.

**Free DNA.** A set of free DNA structures were built using DeepDNAshape [57] (prediction of shape parameters from sequence) and X3DNA-rebuild [58] (structure building based on predicted shape parameters). This set contained all possible DNA hexamers (sampled 4 times each) with random left and right flank added (2-4 base pairs).

| **Dataset** | **PDB** | **AFDB Dist.** | **Interdomain Dist.** | **DNA Dist.** | **DNA interfaces** | **Free DNA** |
|---|---|---|---|---|---|---|
| Approx. example count | $6.95 \times 10^6$ | $7.60 \times 10^6$ | $5.85 \times 10^5$ | $3.83 \times 10^4$ | $2.71 \times 10^4$ | $1.64 \times 10^4$ |
| Spatial crop | 0.75/1.0* | 0.75 | 1.0 | 0.0 | 0.0 | 0.75 |
| Contiguous crop | 0.25/0.0* | 0.25 | 0.0 | 0.4 | 0.4 | 0.25 |
| DNA Contact crop† | 0.0 | 0.0 | 0.0 | 0.6 | 0.6 | 0.0 |
| PPI full binder crop ‡ | 0.75 | 0.0 | 0.75 | 0.0 | 0.0 | 0.0 |
| Sample prob. (stage 1) | 0.2 | 0.7 | 0.1 | 0.0 | 0.0 | 0.0 |
| Sample prob. (stage 2) | 0.2 | 0.5 | 0.1 | 0.14 | 0.04 | 0.01 |

**Table S3:** Probability configuration for different training conditions.

∗ Probabilities for pn_units/interfaces examples, respectively.

†Probabilities given within the subset of examples featurized using the DNA training condition.

‡ Probabilities given within the subset of examples featurized using the PPI training condition.

#### 2 RFdiffusion3 Architecture and Inference

In the following sections, we describe the architecture of RFdiffusion3 in detail. RFdiffusion3 adopts many shared components from AlphaFold3. Features are processed via two parrallel tracks, atom- and token level.

We inherit the use of LinearNoBias, Transition, and ConditionedTransition blocks. For adaptive layer norms and other layer norms we replace all LayerNorm blocks with the simpler RMSNorms in favour of memory efficiency.

##### 2.1 Main inference loop

**Algorithm 1** Inference diffusion loop.

**def** SampleDiffusion($\mathbf{x}_l^{\mathbf{init}}$, $\{\mathbf{f}^*\}$, $\{\sigma_\tau\}_1^T$, $\gamma_0 = 0.6$, $\gamma_{\min} = 1.0$, $\lambda = 1.003$, $\eta = 1.5$):

*# Token and atom embedding*

$(\mathbf{s}_{\text{init}\_i}, \mathbf{z}_{\text{init}\_ij}) \leftarrow$ TokenInitializer($\{\mathbf{f}^*\}$)

$(\mathbf{q}_{\text{init}\_l}, \mathbf{c}_l, \mathbf{p}_{lm}) \leftarrow$ AtomInitializer($\{\mathbf{f}^*\}, \mathbf{s}_{\text{init}\_i}, \mathbf{z}_{\text{init}\_ij}$)

*# Initialize structure*

**for** $\sigma_\tau \in [\sigma_1, \ldots, \sigma_T]$ **do**

*# Modulation of ODE/SDE*

$\gamma \leftarrow \gamma_0$ **if** $\sigma_\tau > \gamma_{\min}$ **else** $0$

$\hat{\sigma} \leftarrow \sigma_{\tau-1}(\gamma + 1)$

*# Noise injection to diffused components*

$\epsilon_l \leftarrow \lambda \sqrt{\hat{\sigma}^2 - \sigma_{\tau-1}^2} \cdot \mathcal{N}(0, I_3) \cdot f^{\text{is\_diffused}}$ $\qquad \epsilon_l \in \mathbb{R}^{L\times 3}$

$\mathbf{x}_l^{\text{noisy}} \leftarrow \mathbf{x}_l + \epsilon_l$

$\{\hat{\mathbf{x}}_0\} \leftarrow$ DiffusionModule($\{\mathbf{x}_l^{\text{noisy}}\}$, $\hat{\sigma}$, $\{f^*\}$, $\mathbf{q}_{\text{init}\_l}$, $\mathbf{c}_l$, $\mathbf{p}_{lm}$, $\mathbf{s}_{\text{init}\_i}$, $\mathbf{z}_{\text{init}\_ij}$ ) $\qquad \hat{\mathbf{x}}_0 \in \mathbb{R}^{L\times 3}$

$d\sigma \leftarrow \sigma_\tau - \hat{\sigma}$

$\mathbf{x}_l \leftarrow \mathbf{x}_l^{\text{noisy}} + \eta \cdot d\sigma \cdot (\mathbf{x}_l - \hat{\mathbf{x}}_0)/\hat{\sigma}$

**end for**

**return** $\{\mathbf{x}_l\}$

##### 2.2 Inference Loop for Symmetric Design

The inference loop for symmetric design follows that of the main inference loop (Algorithm 1), but contains an additional step symmetrizing the coordinates output from the diffusion module. Importantly, this symmetrization step can be stopped prematurely, with the default value stopping symmetrization after 90% of the steps have been run.

**Algorithm 2** Symmetric inference diffusion loop.

**def** SampleSymmetricDiffusion($\mathbf{x}_l^{\mathbf{init}}$, $\{\mathbf{f}^*\}$, $\{\sigma_\tau\}_1^T$, $\gamma_0 = 0.6$, $\gamma_{\min} = 1.0$, $\lambda = 1.003$, $\eta = 1.5$, $\text{mask}_{\text{asu}}$, $\mathbf{R}=\{R_n\}_1^{\text{N}}$, $\sigma_{sym} = 0.9$ ):

*# Token and atom embedding*

$(\mathbf{s}_{\text{init}\_i}, \mathbf{z}_{\text{init}\_ij}) \leftarrow \text{TokenInitializer}(\{\mathbf{f}^*\})$

$(\mathbf{q}_{\text{init}\_l}, \mathbf{c}_l, \mathbf{p}_{lm}) \leftarrow \text{AtomInitializer}(\{\mathbf{f}^*\}, \mathbf{s}_{\text{init}\_i}, \mathbf{z}_{\text{init}\_ij})$

*# Initialize structure*

**for** $\sigma_\tau \in [\sigma_1, \ldots, \sigma_T]$ **do**

*# Modulation of ODE/SDE*

$\quad \gamma \leftarrow \gamma_0$ **if** $\sigma_\tau > \gamma_{\min}$ **else** $0$

$\quad \hat{\sigma} \leftarrow \sigma_{\tau-1}(\gamma + 1)$

*# Noise injection to diffused components*

$\quad \epsilon_l \leftarrow \lambda \sqrt{\hat{\sigma}^2 - \sigma_{\tau-1}^2} \cdot \mathcal{N}(0, I_3) \cdot f^{\text{is\_diffused}}$ $\qquad \epsilon_l \in \mathbb{R}^{L \times 3}$

$\quad \mathbf{x}_l^{\text{noisy}} \leftarrow \mathbf{x}_l + \epsilon_l$

$\quad \{\hat{\mathbf{x}}_0\} \leftarrow \text{DiffusionModule}(\{\mathbf{x}_l^{\text{noisy}}\}, \hat{\sigma}, \{f^*\}, \mathbf{q}_{\text{init}\_l}, \mathbf{c}_l, \mathbf{p}_{lm}, \mathbf{s}_{\text{init}\_i}, \mathbf{z}_{\text{init}\_ij})$ $\qquad \hat{\mathbf{x}}_0 \in \mathbb{R}^{L \times 3}$

*# Symmetrize using the denoised asymmetric unit*

$\quad \hat{\mathbf{x}}_0^{asu} \leftarrow \hat{\mathbf{x}}_0[\text{mask}_{\text{asu}}]$

$\quad \{\hat{\mathbf{x}}_0\} \leftarrow \{R_1 * \hat{\mathbf{x}}_0^{asu}, \ldots, R_N * \hat{\mathbf{x}}_0^{asu}\}$ **if** $(\sigma_\tau / \sigma_T) \geq \sigma_{sym}$

$\quad d\sigma \leftarrow \sigma_\tau - \hat{\sigma}$

$\quad \mathbf{x}_l \leftarrow \mathbf{x}_l^{\text{noisy}} + \eta \cdot d\sigma \cdot (\mathbf{x}_l - \hat{\mathbf{x}}_0)/\hat{\sigma}$

**end for**

**return** $\{\mathbf{x}_l\}$

#### 2.3 Token Initializers

RFdiffusion3 embeds conditioning via both the atom-wise and token-wise tracks. Table S4 summarizes the different conditioning inputs to RFdiffusion3.

| **Feature** | **Dimensions** |
| --- | --- |
| *Token-level features* | |
| ref_motif_token_type | 3 |
| restype | 32 |
| ref_plddt | 1 |
| is_non_loopy | 1 |
| *Atom-level features* | |
| ref_atom_name_chars | 256 |
| ref_element | 128 |
| ref_charge | 1 |
| ref_mask | 1 |
| ref_is_motif_atom_with_fixed_coord | 1 |
| ref_is_motif_atom_unindexed | 1 |
| has_zero_occupancy | 1 |
| ref_pos | 3 |
| ref_atomwise_rasa | 3 |
| active_donor | 1 |
| active_acceptor | 1 |
| is_atom_level_hotspot | 1 |

**Table S4:** Summary of 1D token-level and atom-level input features with their respective dimensionalities.

Token-level features are embedded and passed through an atom-level initializer. The output latent features $\mathbf{c}_l$, $\mathbf{p}_{lm}$, $\mathbf{s}_{\text{init}\_i}$, $\mathbf{z}_{\text{init}\_ij}$ are fed to subsequent blocks as conditioning information after every diffusion step.

**Algorithm 3** Token initializer. Initializes token-level representations from input features and pair encodings. Includes a transformer stack for adding positional information.

**def** TokenInitializer($\{\mathbf{f}^*\}$):

$\mathbf{s}_i \leftarrow$ OneDFeatureEmbedder($\{\mathbf{f}^*\}, I$) $\quad\quad \mathbf{s} \in \mathbb{R}^{I \times c_s}$

$\mathbf{s}_i \leftarrow \mathbf{s}_i$ + Transition($\mathbf{s}_i$)

$\mathbf{s}_i \leftarrow$ Downcast(OneDFeatureEmbedder($\{\mathbf{f}^*\}$, $L$), $\mathbf{s}_i$, $f^{\text{atom\_to\_token\_map}}$)

$\mathbf{s}_i \leftarrow \mathbf{s}_i$ + Transition($\mathbf{s}_i$)

$\mathbf{s}_i \leftarrow$ LinearNoBias(RMSNorm($\mathbf{s}_i$))

*# Initialize pair features with positional encoding, bond graph and reference conformer*

$\mathbf{z}_{ij} \leftarrow$ LinearNoBias($\mathbf{s}_i$) + LinearNoBias($\mathbf{s}_j$) $\quad\quad \mathbf{z} \in \mathbb{R}^{I \times I \times c_z}$

$\mathbf{z}_{ij} \leftarrow \mathbf{z}_{ij}$ + RelativePositionEncodingWithIndexRemoval($\{\mathbf{f}^*\}$)

$\mathbf{z}_{ij} \leftarrow \mathbf{z}_{ij}$ + outer(LinearNoBias($f^{\text{token\_bonds}}$))

$\mathbf{z}_{ij} \leftarrow \mathbf{z}_{ij}$ + PositionPairDistEmbedder($f^{\texttt{ref\_pos, is\_ca}}$, mask)

*# Transformer-based position mixing*

**for** $b = [1, 2]$ **do**

  $\mathbf{s}_i, \mathbf{z}_{ij} \leftarrow$ TransformerBlock($\mathbf{s}_i$, $\mathbf{z}_{ij}$)

**end for**

$\mathbf{z}_{ij} \leftarrow$ concat($\mathbf{z}_{ij}$, RelativePositionEncodingWithIndexRemoval($\{\mathbf{f}^*\}$) $\quad\quad$ [Fig. S11]

$\mathbf{z}_{ij} \leftarrow$ LinearNoBias(RMSNorm($\mathbf{z}_{ij}$))

**for** $b = [1, 2]$ **do**

  $\mathbf{z}_{ij} \leftarrow$ RMSNorm($\mathbf{z}_{ij}$)

  $\mathbf{z}_{ij} \leftarrow \mathbf{z}_{ij}$ + Transition($\mathbf{z}_{ij}$)

**end for**

**return** $\{\mathbf{s}_i\}, \{\mathbf{z}_{ij}\}$

**Algorithm 4** Atom initializer. Embeds atom-level representations, integrates pairwise and sequence-local features, and applies optional 2D conditioning.

**def** AtomInitializer($\{\mathbf{f}^*\}, \{\mathbf{s}_i\}, \{\mathbf{z}_{ij}\}$):

$\{\mathbf{q}_{\text{init\_}l}\} \leftarrow \text{OneDFeatureEmbedder}(\{\mathbf{f}^*\}, L)$

$\mathbf{c}_l \leftarrow \mathbf{q}_{\text{init\_}l} + \text{LinearNoBias}(\text{RMSNorm}(\mathbf{s}_{\text{tok\_idx}(l)}))$

*# Embed motif positions and ref coordinates*

$\mathbf{p}_{lm} \leftarrow \text{SinusoidalDistEmbed}(f^{\text{motif\_pos}}, f^{\text{is\_motif\_fixed\_coord}})$

$\mathbf{p}_{lm} \leftarrow \mathbf{p}_{lm} + \text{PositionPairDistEmbedder}(f^{\texttt{ref\_pos}})$

*# Project atom single and pair features*

$\mathbf{p}_{lm} \leftarrow \mathbf{p}_{lm} + \text{LinearNoBias}(\text{ReLU}(c_{\text{tok\_idx(l)}})) + \text{LinearNoBias}(\text{ReLU}(c_{\text{tok\_idx(m)}}))$

$\mathbf{p}_{lm} \leftarrow \mathbf{p}_{lm} + \text{LinearNoBias}(\text{RMSNorm}(z_{\text{tok\_idx(l)},\text{tok\_idx(m)}}))$

$\mathbf{p}_{lm} \leftarrow \mathbf{p}_{lm} + \text{LinearNoBias}(\text{ReLU}(\text{LinearNoBias}(\text{ReLU}(\text{LinearNoBias}(\text{ReLU}(\mathbf{p}_{lm}))))))$

**return** $\{\mathbf{q}_{\text{init\_}l}\}, \{\mathbf{c}_l\}, \{\mathbf{p}_{lm}\}$

#### 2.4 DiffusionModule

The diffusion module of RFdiffusion3 resembles that of a UNet-style architecture [16], similar to AlphaFold3. The diffusion module serves to process the noisy coordinates alongside the initialized conditional features to predict the denoised structure. In contrast to AF3, we replace mean pooling of the diffusion module with a learnable cross attention (Fig. S12). Similar to previous work [12, 1, 3, 2], we use recycling of the central-atom distogram; notably, we do not recycle atom-level features. Prioritizing efficient computation we do not use any triangle multiplicative updates within RFdiffusion3. In algorithm Algorithm 5, we summarize the main forward pass of the diffusion module.

**Algorithm 5** Diffusion forward pass with recycling. Computes denoised positions from noisy coordinates and initial token/atom features.

**def** DiffusionModule($\{\mathbf{x}_l^{\text{noisy}}\}, \sigma, \{\mathbf{f}^*\}, \{\mathbf{q}_l^{\text{init}}\}, \{\mathbf{c}_l\}, \{\mathbf{p}_{lm}\}, \{\mathbf{s}_i\}, \{\mathbf{z}_{ij}\}$):

*# Scale positions to dimensionless vectors with approximately unit variance.*

$r_\ell^{\text{noisy}} \leftarrow \left(\frac{1}{1+(\sigma/\sigma_{\text{data}})^2}\right) \cdot \mathbf{x}_l^{\text{noisy}}$

*# Pool to sequence level and downcast*

$\mathbf{a}_i \leftarrow \underset{\text{tok\_idx}(l)}{\text{mean}} \text{LinearNoBias}(r_l^{\text{noisy}})$

$\mathbf{s}_i \leftarrow \text{Downcast}(\mathbf{c}_l, \mathbf{s}_i)$

*# Add batch/time embeddings*

$\mathbf{q}_l \leftarrow \mathbf{q}_l^{\text{init}} + \text{LinearNoBias}(r_l^{\text{noisy}})$

$\mathbf{c}_l \leftarrow \mathbf{c}_l + \text{ProcessTime}(\sigma \cdot f^{\text{is\_unfixed}})$

$\mathbf{s}_i \leftarrow \mathbf{s}_i + \text{ProcessTime}(\sigma_l \cdot f^{\text{is\_unfixed}})$

$\mathbf{c}_l \leftarrow \mathbf{c}_l + \text{LinearNoBias}(\text{RMSNorm}(\mathbf{c}_l))$

*# Local-atom transformer + pooling*

$\mathbf{q}_l \leftarrow \text{SparseTransformer}(\mathbf{q}_l,\ \mathbf{c}_l,\ \mathbf{p}_{lm},\ \mathbf{x}_l^{\text{noisy}},\ \emptyset,\ n_{\text{block}} = 3)$

$\mathbf{a}_i \leftarrow \text{Downcast}(\mathbf{q}_l, \mathbf{a}_i, \mathbf{s}_i)$

*# Process with recycling*

$\{\hat{\mathbf{x}}_0\}, \{\mathbf{s}_i^{\text{init}}\}, \{\mathbf{z}_{ij}^{\text{init}}\} \leftarrow (\emptyset,\ \mathbf{s}_i,\ \mathbf{z}_{ij})$ $\quad \hat{\mathbf{x}}_0 \in \mathbb{R}^{L\times 3}$

**for** $n_{\text{recycle}} \in [1, 2]$ **do**

*# Embed noise scale and recylced distogram*

  $\mathbf{s}_i, \mathbf{z}_{ij} \leftarrow \text{DiffusionTokenEncoder}(\mathbf{s}_i^{\text{init}}, \mathbf{z}_{ij}^{\text{init}}, \{\mathbf{f}^*\}, \mathbf{x}_l^{\text{noisy}}, \hat{\mathbf{x}}_0, n_{\text{block}} = 2)$

*# Sparse attention at the token level*

  $\mathbf{a}_i \leftarrow \text{SparseTransformer}(\mathbf{a}_i,\ \mathbf{s}_i,\ \mathbf{z}_{ij},\ \mathbf{x}_l^{\text{noisy}},\ \emptyset,\ n_{\text{block}} = 18)$

*# Up-projection and decode to structure*

  $\mathbf{q}_l \leftarrow \text{SparseTransformer}(\mathbf{q}_l,\ \mathbf{c}_l,\ \mathbf{p}_{lm},\ \mathbf{x}_l^{\text{noisy}},\ \mathbf{a}_i,\ n_{\text{block}} = 3)$

  $\mathbf{r}_l^{\text{update}} \leftarrow \text{LinearNoBias}(\mathbf{q}_l)$ $\quad \mathbf{r}_l^{\text{update}} \in \mathbb{R}^{L\times 3}$

  $\hat{\mathbf{x}}_0 \leftarrow \left(\frac{1}{1+(\sigma/\sigma_{\text{data}})^2}\right) \cdot \mathbf{x}_l^{\text{noisy}} + \left(\frac{\sigma/\sigma_{\text{data}}}{\sqrt{1+(\sigma/\sigma_{\text{data}})^2}}\right) \cdot \mathbf{r}_l^{\text{update}}$

**end for**

**return** $\{\hat{\mathbf{x}}_0\}$

###### 2.4.1 Atom Encoder, Token Transformer and Decoder

An atom encoder operates on all atom features and coordinates from the developing noise cloud. The generated atomic features are pooled into token features and fed through a token-wise transformer. Finally, the features are expanded back to the atom level progressively while an atom decoder operates on the output token features. There are three blocks of Upcast (Algorithm 10) interweaved in the SparseTransformer (Algorithm 6) for the readout. The final atom features are then used to predict the updates to the all-atom coordinates.

###### 2.4.2 Sequence- and structure-local sparse attention

**SL2 attention.** Inspired by the efficient sequence-local atom attention of AF3, as well as sparse attention variants for diffusion [66], we use a sequence-and-structure local atom attention variant within RFdiffusion3 (*SL2* attention). In particular, we first use sequence-local attention consisting of a tunable $n$-adjacent tokens, and then pad the remaining number of keys based on positions local in the noise. During the second forward pass with recycling, we use the models' own prediction to determine the indices rather than relying on atomic positions within the noise. In Fig. S12, we show an example attention mask with these components.

**Attention keys selection.** First, sequence neighbors are selected (using $n$-adjacent tokens on either side of the given token). We ensure all atoms in neighboring tokens are included (14 for canonical residues); that is, we do not partially attend to a given token while expanding with sequence-local attention. Second, the remaining attention keys are selected by using a distance matrix of the distances in the noise (or the models' own prediction, for the second forward pass with recycling) and selecting the $k$ atoms that are closest to each other atom in Euclidean space until each atom meets its key budget.

At the token level, it is not strictly necessary to use sparse attention; however, we found that sparse attention regularizes the model to prevent overfitting and drastically accelerates inference at larger token counts. For token attention, we use $n^{\text{token}}_{\text{sequence\_neighbours}} = 32$, and $n^{\text{token}}_{\text{keys}} = 128$. For atom attention, we use $n^{\text{atom}}_{\text{sequence\_neighbours}} = 32$ and $n^{\text{atom}}_{\text{keys}} = 128$.

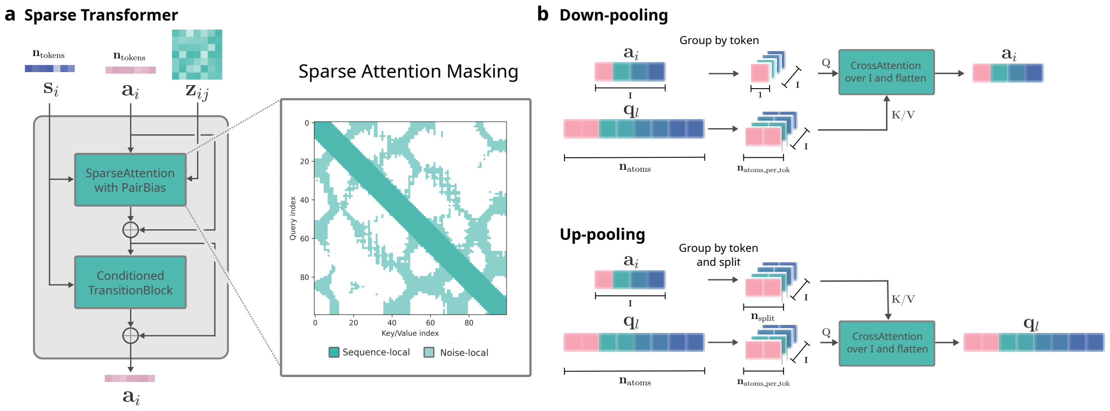

**Fig. S12: RFD3 Architecture components.** **a**, Transformer blocks in RFdiffusion3 follow a modern architecture with standard attention and conditioned transition blocks. Additionally, we use a sparse attention variant of the standard attention architecture. **b**, Atom-level features are processed via a UNet-type architecture (Fig. 1).

**Algorithm 6** Local token transformer. Computes local attention over tokens. Attention indices are precomputed based on local sequence and structure neighborhoods using SL2 attention.

**def** SparseTransformer($\mathbf{q}_l$, $\mathbf{c}_l$, $\mathbf{p}_{lm}$, $\{\mathbf{f}^*\}$, $\mathbf{x}_l^{\text{noisy}}$, $\mathbf{c}_{\text{skip}}$, $n_{\text{block}}$):

*# Compute local attention indices for each token*

indices $\leftarrow$ create_attention_indices($\hat{\mathbf{x}}_l^{\text{self}}$, $\{\mathbf{f}^*\}$) **if** $\mathbf{x}_l^{\text{noisy}}$ **else** null — Figure S12a

**for** _ in $\{1, 2, \ldots, n_{\text{block}}\}$ **do**

  **if** $\{\mathbf{c}_{\text{skip}}\} \neq \emptyset$ **then**

    $\mathbf{q}_l \leftarrow$ Upcast($\mathbf{q}_l, \mathbf{c}_{\text{skip}}$)

  **end if**

  $\mathbf{q}_l \leftarrow \mathbf{q}_l +$ SparseAttentionPairBias($\mathbf{q}_l$, $\mathbf{c}_l$, $\mathbf{p}_{lm}$, indices) — $\mathbf{q}_l \in \mathbb{R}^{B \times L \times c_{\text{atom}}}$ or $\mathbb{R}^{B \times L \times c_{\text{token}}}$

  $\mathbf{q}_l \leftarrow \mathbf{q}_l +$ ConditionedTransitionBlock($\mathbf{q}_l$, $\mathbf{c}_l$)

**end for**

**return** $q_i$

**Algorithm 7** For processing pair features into the single track, we use a trimmed-down pairformer module which is equivalent to full attention.

**def** TransformerBlock($\mathbf{s}_i$, $\mathbf{z}_{ij}$):

$\mathbf{s}_i \leftarrow \mathbf{s}_i +$ SparseAttentionPairBias($\mathbf{s}_i$, $\emptyset$, $\mathbf{z}_{ij}$, $\emptyset$, full=True) — $\mathbf{s}_i \in \mathbb{R}^{I \times c_{\text{token}}}$

$\mathbf{s}_i \leftarrow \mathbf{s}_i +$ Transition($\mathbf{s}_i$)

**return** $\{\mathbf{s}_i, \mathbf{z}_{ij}\}$

**Algorithm 8** Sparse attention with pair bias. For tokens, $q_l \rightarrow a_i, c_l \rightarrow s_i, p_{ll} \rightarrow z_{ii}$. sparse_pairbias_attention is a simple implementation of attention pair bias using the attention indices provided, computing either the full attention matrix or attended indices only.

**def** SparseAttentionPairBias($\{\mathbf{q}_l\}$, $\{\mathbf{c}_l\}$, $\{\mathbf{p}_{lm}\}$, indices, full=False):

**if** $\mathbf{c}_l$ exists **then**

  $\mathbf{q}_l \leftarrow \mathrm{AdaLN}(\mathbf{q}_l, \mathbf{c}_l)$

**else**

  $\mathbf{q}_l \leftarrow \mathrm{RMSNorm}(\mathbf{q}_l)$

**end if**

*# Project to queries, keys, values, gate, and bias* $\quad q, k, v, g \in \mathbb{R}^{H \times L \times L \times c_{\text{head}}}$

$q \leftarrow \mathrm{LinearNoBias}(\mathbf{q}_l)$

$k, v \leftarrow \mathrm{RMSNorm}(\mathrm{LinearNoBias}(\mathbf{q}_l))$

$g \leftarrow \mathrm{Sigmoid}(\mathrm{LinearNoBias}(\mathbf{q}_l))$

$b \leftarrow \mathrm{LinearNoBias}(\mathbf{p}_{lm})$ $\quad h \in \mathbb{R}^{L \times L \times H}$

*# Sparse attention with pairwise bias*

$\alpha \leftarrow$ sparse_pairbias_attention($q$, $k$, $v$, $b$, indices, $g$, full) $\quad \alpha \in \mathbb{R}^{L \times L \times c_{\text{head}}}$

$o \leftarrow \mathrm{LinearNoBias}(\alpha)$ $\quad o \in \mathbb{R}^{L \times L \times c_{\text{atom}}}$

*# Optionally gate with conditioning signal*

**if** $\mathbf{c}_l$ exists **then**

  $o \leftarrow \mathrm{sigmoid}(\mathrm{Linear}(\mathbf{c}_l, \text{biasinit}{=}{-2.0})) \odot o$

**end if**

**return** $o$

#### 2.5 Cross attention pooling and broadcast

#### 2.6 Embedders

**Algorithm 9** Downcast

**def** Downcast($q_l$, $a_i$, $s_i$):

*# Group atoms by corresponding token id*

$q_{ia} = \text{reshape}(q_l)$ $\quad q_{ia} \in \mathbb{R}^{c \times a}$

$q_l = q_l + \text{flatten}(\text{GatedCrossAttention}(Q{=}a_i,\ KV{=}q_{ia}))$

**if** $\{s_i\} \neq 0$ **then**

$\quad a_i \mathrel{+}= \text{LinearNoBias}(s_i)$

**end if**

**return** $a_i$

**Algorithm 10** Upcast

**def** Upcast($\mathbf{q}_l$, $\mathbf{a}_i$):

*# Split tokens and group atoms by corresponding token id*

$\mathbf{a}_i^{\text{split}} = \text{reshape}(\mathbf{a}_i)$ $\quad \mathbf{a}_i^{\text{split}} \in \mathbb{R}^{I \times \text{n}_{\text{split}} \times c'}$

$\mathbf{q}_{ia} = \text{reshape}(\mathbf{q}_l)$ $\quad \mathbf{q}_{ia} \in \mathbb{R}^{I \times a \times c}$

$\mathbf{q}_l = \mathbf{q}_l + \text{flatten}(\text{GatedCrossAttention}(Q{=}\mathbf{q}_{ia},\ KV{=}\mathbf{a}_i^{\text{split}}))$

**return** $\mathbf{q}_l$

**Algorithm 11** Gated cross attention for upcast and downcasting

**def** GatedCrossAttention($q$, $kv$, valid_mask=None):

*# Normalize input representations*

$q \leftarrow \text{RMSNorm}(q)$

$kv \leftarrow \text{RMSNorm}(kv)$

*# Project to query, key, value, gate*

$q \leftarrow \text{LinearNoBias}(q)$

$k \leftarrow \text{RMSNorm}(\text{LinearNoBias}(kv))$

$v \leftarrow \text{RMSNorm}(\text{LinearNoBias}(kv))$

$g \leftarrow \text{Sigmoid}(\text{LinearNoBias}(q))$

*# Scaled dot-product attention*

$\alpha \leftarrow \frac{1}{\sqrt{c}} \cdot \sum_k q_{ic} \cdot k_{jc}^{\top}$ (contract over channel dim $c$)

*# Apply valid mask and remove invalid queries where no keys are valid*

**if** valid_mask exists **then**

  $\alpha \leftarrow \alpha$.masked_fill($\sim$ valid_mask, $-\infty$)

  $\alpha[\sim$ valid_mask.any$(-2)] \leftarrow 0$

**end if**

*# Softmax and value aggregation*

$\alpha \leftarrow \text{Softmax}(\alpha, \text{dim} = k)$

$o_{qc} \leftarrow \sum_k \alpha_{qk} \cdot v_{kc}$

*# Merge heads and project*

$o \leftarrow \text{Linear}(o \cdot g)$

**return** $o$

#### 2.7 Classifier Free Guidance

Classifier-free guidance is a technique originally developed in diffusion models to enhance conditional generation by amplifying the influence of conditioning signals during inference [20]. RFdiffusion3 is trained to handle both conditional (with design specifications) and unconditional (without design specifications) generations. During inference, the guided prediction is computed as:

$$\tilde{\epsilon}_\theta(\mathbf{x}_t|\mathbf{c}) = \epsilon_\theta(\mathbf{x}_t) + (\omega - 1)(\epsilon_\theta(\mathbf{x}_t|\mathbf{c}) - \epsilon_\theta(\mathbf{x}_t)) \tag{5}$$

where $\tilde{\epsilon}_\theta(\mathbf{x}_t|\mathbf{c})$ is the step taken under CFG, $\epsilon_\theta(\mathbf{x}_t|\mathbf{c})$ is the conditional step, and $\epsilon_\theta(\mathbf{x}_t)$ is the unconditional step, with $\omega \geq 1$ is the guidance scale (CFG-scale) that controls the strength of conditioning adherence. A higher CFG-scale prioritizes adherence to specifications, commonly at the expense of sample diversity. Classifier-free guidance has been employed in prior protein design frameworks to guide models toward high-level structural features, such as fold class annotations. Geffner et al. [12] utilized CFG to steer the diffusion-based models process toward specific fold classes. Unlike previous approaches, which used classifier-free guidance for global conditioning, we for the first time demonstrate atomic-level CFG.

**Algorithm 12** The diffusion token encoder serves to condition the token forward pass with recycling information. Distogram features are bucketized according to a minimum distance of 1 Åand a maximum of 30 Å, with 65 linearly-spaced bins.

**def** DiffusionTokenEncoder($\mathbf{s}_i^{\text{init}}$, $\mathbf{z}_{ij}^{\text{init}}$, $\{\mathbf{f}^*\}$, $\mathbf{x}_l^{\text{noisy}}$, $\hat{\mathbf{x}}_l^{\text{self}} = \emptyset$, $n_{\text{block}} = 2$):

**for** _ in $\{1, 2\}$ **do**

  $\mathbf{s}_i \leftarrow \mathbf{s}_i$ + Transition($\mathbf{s}_i$)

**end for**

$\mathbf{z}_{ij}^{\text{concat}} \leftarrow$ concat($\mathbf{z}_{ij}$, distogram($\mathbf{x}_l^{\text{noisy}}$), distogram($\hat{\mathbf{x}}_l^{\text{self}}$))

$\mathbf{z}_{ij} \leftarrow$ LinearNoBias(RMSNorm($\mathbf{z}_{ij}$))

**for** _ in $\{1, 2\}$ **do**

  $\mathbf{z}_{ij} \leftarrow \mathbf{z}_{ij}$ + Transition($\mathbf{z}_{ij}$, $n = 2$)

**end for**

*# Mix into conditioning track with Pairformer*

**for** _ in $\{1, 2, \ldots, n_{\text{block}}\}$ **do**

  $\{\mathbf{s}_i, \mathbf{z}_{ij}\} \leftarrow$ TransformerBlock($\mathbf{s}_i$, $\mathbf{z}_{ij}$)

**end for**

**return** $\{\mathbf{s}_i, \mathbf{z}_{ij}\}$

**Algorithm 13** Sinusoidal Distance Embedding. The **mask** defines the valid off-diagonals.

**def** SinusoidalDistEmbed($\{\mathbf{x}_l\}, \{\mathbf{mask}\}$)

*# Calculate pairwise distances and perform sinusoidal embedding.*

$\mathbf{d}_{lm} \leftarrow \|\mathbf{x}_l - \mathbf{x}_m\|_2$

$\omega_k \leftarrow 10000^{-k/n_{\text{freq}}}$

$\boldsymbol{\theta}_{lmk} \leftarrow D_{lm} \cdot \omega_k$

$\mathbf{e}_{lm} \leftarrow [\sin(\boldsymbol{\theta}_{lmk}) \parallel \cos(\boldsymbol{\theta}_{lmk})]$

*# Apply and embed masks.*

$\mathbf{p}_{lm} \leftarrow \text{LinearNoBias}(\mathbf{e}_{lm})$

$\mathbf{p}_{lm} \leftarrow \mathbf{p}_{lm} \cdot \mathbf{mask}_{lm}$

$\mathbf{p}_{lm} \leftarrow \mathbf{p}_{lm} + \text{LinearNoBias}(\mathbf{mask}_{lm}) \cdot \mathbf{mask}_{lm}$

**return** $\{\mathbf{p}_{lm}\}$

**Algorithm 14** Position Pair Distance Embedding.

**def** PositionPairDistEmbedder($\{\mathbf{x}_l\}, \{\mathbf{mask}\}$)

*# Calculate pairwise distances.*

$\mathbf{d}_{lm} \leftarrow \|\mathbf{x}_l - \mathbf{x}_m\|_2$

*# Encode the inverse pairwise distance.*

$\mathbf{inv}_{lm} \leftarrow 1/(1 + \max(\|\mathbf{d}_{lm}\|^2, 10^{-6}))$

*# Embed the mask.*

$\mathbf{p}_{lm} \leftarrow \text{Linear}(\mathbf{inv}_{lm}) \cdot \mathbf{mask}_{lm}$

$\mathbf{p}_{lm} \leftarrow \mathbf{p}_{lm} + \text{Linear}_{\text{mask}}(\mathbf{mask}_{lm}) \cdot \mathbf{mask}_{lm}$

**return** $\{\mathbf{p}_{lm}\}$

**Algorithm 15** One-dimension Feature Embedder for both token- and atom-level features.

**def** OneDFeatureEmbedder($\{\mathbf{f}^*\}$)

*# Embed each 1D feature and sum results.*

$\mathbf{e} \leftarrow \mathbf{0}$

**for** each feature $k$ with channels $n_k$ **do**

    **if** $n_k$ exists **then**

        $\mathbf{e}_k \leftarrow LinearNoBias(\mathbf{f}^k)$

        $\mathbf{e} \leftarrow \mathbf{e} + \mathbf{e}_k$

    **end if**

**end for**

**return** $\mathbf{e}$

**Algorithm 16** FourierEmbedding

**def** FourierEmbedding($t_l$):

*# Initialize parameters*

$w, b \sim \mathcal{N}(0, 1)$

*# Compute Fourier features*

$\mathbf{o} \leftarrow \cos(2\pi \cdot (t_l \cdot w + b))$

**return** $\{\mathbf{o}\}$

**Algorithm 17** Processing of noise conditioning features.

**def** ProcessTime($t_l$):

*# Embed time with Fourier and project*

$\mathbf{c}_l \leftarrow \mathrm{LinearNoBias}(\mathrm{RMSNorm}(\mathrm{FourierEmbedding}(\frac{1}{4}\log(\frac{\max(t_L, 1e-20)}{\sigma_{\text{data}}}))))$

$\mathbf{c}_l \leftarrow \mathbf{c}_l \odot (t_l > 0)$

**return** $\{\mathbf{c}_l\}$

### 3 *In silico* Evaluation

#### 3.1 Unconditional evaluation

Unconditional backbones of length 100, 150 and 250 residues were sampled from the model (32 of each) at `step_scale` 1.0, 1.25 and 1.5. The resulting backbones were redesigned 8 times using ProteinMPNN and refolded with AlphaFold3 [11, 25]. Backbones with N, C$\alpha$, C RMSD $\leq$ 1.5 Å when compared to the design were considered an *in silico* success. Backbones were then searched for similar homologs in the PDB using Foldseek and clustered using pairwise TM-align.

Furthermore, we evaluated the computational efficiency of unconditional monomer generation using RFD3 (Table S5). For lengths of 50, 100, 200, and 400 residues, we generated 16 designs each using RFdiffusion1, RFdiffusion2, and RFdiffusion3 with one NVIDIA a6000 GPU, 1 CPU core, and 32 GB of RAM. We then computed the average time required to generate a single design. RFdiffusion3 was run with batch size 16 and 200 denoising steps, whereas RFdiffusion1, RFdiffusion-AA and RFdiffusion2 were run with 50 steps. These previous models do not support a batch size greater than 1.

| Num tokens / Model(s/design) | RFD3 | RFD1 | RFD2 |
|---|---|---|---|
| **50** | 2.53 | 24.26 | 41.03 |
| **100** | 4.18 | 24.98 | 42.64 |
| **200** | 8.97 | 43.88 | 71.93 |
| **400** | 23.01 | 153.68 | 234.34 |

**Table S5:** Efficiency of protein monomer generation using RFdiffusion3.

##### 3.2 Protein-protein interaction benchmark

Five protein targets were chosen for the protein-protein interaction benchmark: three that have been benchmarked in other works (PD-L1, InsulinR, IL-7Ra) and two that have not (Tie2, IL-2Ra). Details of the setup for each target are specified in Table S6. Length ranges for previously-benchmarked targets were chosen to align with [8], while for the new targets we chose a larger length range that more accurately reflects our typical design campaigns. The hotspots, approximate center-of-mass of the binder, and the target structures were provided as conditioning to RFD3. All other hyperparameters were set to the default values. The center-of-mass conditioning was selected as follows: the mean position of all target atoms within 12 Å of any hotspot atom was computed, and a vector was constructed from this point to the mean position of the hotspot atoms themselves. This vector was then extended an additional 10 Å from the mean position of the hotspots to obtain the center-of-mass value used to initialize the noisy coordinates.

For each target, we generated 400 unique backbones with RFD3 and 400 with RFD1, then generated 4 sequences each with ProteinMPNN. When running RFD1, we provide hotspots for each residue that contains any of the atom-level hotspots given to RFD3. Following the finding in [1] that setting the noise scale to 0 improves designability of RFD1 outputs, we set `noise_scale_ca` = 0 and `noise_scale_frame` = 0 when benchmarking RFD1. In Fig. 3, we report both the total number of backbones and the total number of backbone clusters with at least one sequence passing the AF3 refolding cutoffs from [8]. Agglomerative clustering is performed at a TM-score threshold of 0.6 using the "complete linkage" criterion, meaning that any given pair of elements in a cluster must have a TM-score above the threshold value. In Fig. S2, we report additional designability metrics: the unclustered pass rate among all sequence-structure pairs, normalized to 100 backbones (with 4 MPNNs) per target; the overall distributions of AF3 minimum inter-chain pAE values; and the same metrics among only designs with >75% helix content. We also report monomer diversity AUC values for each target, computed using the "cluster fraction" (number of clusters divided by number of examples) for TM-score clustering thresholds spaced at intervals of 0.1 between 0 and 1, exclusive. Last, we include analogous AUC curves using a new metric, the "dock diversity", defined as the target-aligned backbone N, C$\alpha$, C RMSD between the 15 residues in the design that are closest to the target by C$\alpha$-C$\alpha$ distance. The cluster thresholds are evenly spaced at intervals of 0.1 Å between 0 and 15, inclusive. As before, the designed backbones are clustered agglomeratively using the "complete linkage" criterion.

| Target | PDB ID | Target chain and residues | Hotspot atoms | Design lengths |
|---|---|---|---|---|
| PD-L1 | 5o45 | A17-131 | A56: CG, OH; A115: CG, SD; A123: CD2, OH | 50-120 |
| InsulinR | 4zxb | E6-155 | E64: CD2, CZ; E88: CG, CZ; E96: CD1, CZ | 40-120 |
| IL-7Ra | 3di3 | B17-209 | B58: CG1, CG2; B80: CG, CD1; B139: CD2, OH | 50-120 |
| Tie2 | 2gy5 | A23-210 | A154: CG1, CG2; A156: CG, CZ; A157: CG, CD A161: CG, CZ; A162: CG2, CD1; A176: CG1, CG2 A178: CG, CD1; A189: CG, CZ; A194: CG2, CD1 | 65-120 |
| IL-2Ra | 1z92 | B0-47,B53-64, B104-165 | B2: CB, CD1; B25: SD, CE; B42: CB, CD1 B43: CD2, OH; B120: CD2, CE1 | 65-120 |

**Table S6:** Details of PPI Benchmark. Following previous work, all residue indices refer to the author-annotated values. The asymmetric unit is used for all structures. All ranges are inclusive, both for target residues and for design lengths. No relaxation step or other preprocessing is performed at any stage in target preparation – the file downloaded from the RCSB PDB is simply cropped to the chains/indices given above and then provided directly to the model.

#### 3.3 DNA binder benchmark

Target DNA were chosen from three PDB entries held-out from the RFD3 training set - 7M5W, 7RTE and 7N5U. In each case, a non-repetitive 10 base pair stretch of DNA was chosen, to control for any potential effect of DNA length on the success rate. For evaluation targeting flexible DNA, only the corresponding DNA sequences were used as input. For evaluation targeting fixed DNA, the DNA atomic coordinates used were from biological assembly no. 1 of these PDB entries.

For each DNA target and setting (fixed vs flexible DNA), we generate 100 designs with length sampled between 50-80 residues. Using LigandMPNN we sample 4 sequences per backbone, then refold the protein-DNA pairs using Alphafold3. We assess quality by first aligning the DNA in the design model and the AF3 prediction, then computing the RMSD between protein $C_\alpha$ atoms in the design model and the AF3 prediction. Using the Biotite [63] implementation of the P-SEA algorithm [67] to assign secondary structure, we remove contiguous residues on the N or C termini that are not assigned as helices or sheets. The purpose of this is to avoid penalizing the network for generating loopy regions in the termini, as these are common in DNA binding proteins but are less likely to refold with low RMSD. For fixed DNA cases, we initialize the center of mass of the diffused protein by computing the center of mass of the phosphate atoms, then adding a 3 Åvector parallel to the direction of the major groove.

#### 3.4 Small-molecule benchmark

To evaluate the effectiveness of RFdiffusion3 in small-molecule binder design, we selected four commonly used targets from the RosettaFold All-Atom [2] and BoltzDesign [42] benchmarks: FAD (PDB ID: 7BKC), OQO (PDB ID: 7V11), IAI (PDB ID: 5SDV), and SAM (PDB ID: 7C7M). We benchmark the model under two ligand settings: fixed ligands and diffused ligands.

In the fixed ligand setting, we use the exact all-atom coordinates from the PDB structure. This allows the ligand to serve as a rigid motif, around which the model is expected to generate a compatible binding scaffold. In this setting, the center of mass of the ligand is used to initialize the noise cloud. In the diffused ligand setting, the model does not receive explicit coordinates.

Instead, we used RDKit to generate the ligand's reference conformer and passed this as a feature to the model, which was then required to co-diffuse the ligand with the protein. This setting leads to greater variability in the resulting ligand conformations. To encourage better binding poses, we applied RASA conditioning to guide the model toward designs that bury the ligand, using classifier-free guidance (CFG) with a scale of 2 to strengthen this conditioning.

We establish four benchmark tasks to evaluate small-molecule binder design:

1. **Designability.** For each ligand target and each ligand setting, we generate 400 designs with length of 150 using RFdiffusion3. For each design, we use LigandMPNN to generate 8 sequences, and then refold the sequence-ligand pairs using Alphafold3. We assess quality using two criteria: (i) protein backbone RMSD (after alignment), and (ii) ligand RMSD after aligning the protein backbones. A design is considered successful if: backbone RMSD $\leq$ 1.5 Å & backbone-aligned ligand RMSD $\leq$ 5 Å & interface min pae $\leq$ 1.5 & ipTM $\geq$ 0.8. Results from Fig. S4f show that RFdiffusion3 achieves a higher in silico success rate across all four ligands.

2. **Diversity.** To quantify structural diversity, we apply TM-align with an agglomerative clustering algorithm to cluster the 400 designs, then report the number of clusters under 10 different TM-score thresholds between 0.0 and 1.0. Results in Fig. S4a show that the fixed-ligand setting produces more diverse structures than the diffused-ligand setting. We hypothesize that in the diffused-ligand case, the model probably has less incentive to explore protein conformational space, since it can instead adjust the ligand conformation when struggling to find a compatible protein.

3. **Binding energy.** In some cases, the model generates plausible helical bundles that are designable according to AlphaFold3 and LigandMPNN, but are physically unrealistic. To better assess true binding, we used Rosetta to compute the $\text{DDG}_{\text{norepack}}$ metric, which estimates the change in energy between bound and unbound states. As shown in S4c, RFdiffusion3 designs exhibit stronger binding, reflected by lower $\Delta\Delta G$ values.

4. **Novelty.** We assess structural novelty by querying each of the 400 designs per target against the PDB using FoldSeek, and report the TM-score of the closest structural neighbor (Fig. S4. Lower TM-scores indicate more novel designs.

To evaluate the conformational quality of diffused ligands, we used RDKit to generate 50 conformers for each ligand target. For each design, we computed the RMSD to all 50 conformers and recorded the minimum RMSD. The structures corresponding to the minimum RMSD among all designs, along with the full RMSD distribution, are shown in Fig. S4d-e.

Additionally, we aimed to evaluate whether the interface residue identities were accurately captured. To define the interface, we selected all residues within 2.5–3 Å of any ligand atom. These interface residues were then fixed, and LigandMPNN was used to design 8 sequences for the remaining regions of each structure. The resulting sequences were refolded using AlphaFold3, and their designability was evaluated using the same metrics described previously. A comparison between these constrained designs and those generated by full-sequence LigandMPNN is presented in Fig. S4f.

##### 3.5 Atomic motif enzyme (AME) benchmark

We use the atomic motif enzyme (AME) benchmark to evaluate RFdiffusion3. The AME benchmark consists of a set of 41 cases for scaffolding tipatoms and ligands.

Pass rate for each case is defined as the fraction of designs in 100 backbones that satisfy two conditions:

1. **Chai-1 motif sidechain recapitulation:** The backbone passes this criterion if any of 5 Chai models (one trunk seed) for any of the 8 LigandMPNN sequences for the backbone has a motif backbone (N, C$\alpha$, C) aligned, motif all-atom RMSD $\leq 1.5$ Å.
2. **No ligand clashes.** A ligand clash is defined as any non-motif backbone atom (N, CA or C) which is within $\leq 1.5$ Å of the closest ligand atom.

For each case, we run 10 batches of diffusion batch size 10. For exact details of the constraints of the 41 different cases, we refer the reader to [3].

##### 3.6 Unconditional Symmetric Backbone Generation

We evaluated the ability of RFdiffusion3 to unconditionally generate homo-oligomers across cyclic and dihedral symmetries (C2-C7, D2). For each case, we generate 100 RFdiffusion3 backbones, each with a 100-residue asymmetric unit (ASU). We then sample 8 sequences per backbone with ProteinMPNN and predict the full protein complexes with five AlphaFold3 models (one model seed). A backbone is considered a success if any of the 8 ProteinMPNN sequences re-folds via AlphaFold3 with a complex C$\alpha$ RMSD of $\leq 2.0$ Å and pTM $\geq 0.5$.

##### 3.7 Symmetric Atom Motif Scaffolding Benchmark

We also evaluated the ability of RFdiffusion3 to design symmetric enzymes (Fig. S6). We use a subset of the AME benchmark to do so:

1. **C2 symmetry.** In the AME benchmark, there are several examples of native enzymes with C2 symmetry. We subselected these examples for the symmetric AME benchmark, excluding any examples where the symmetric frames deviate from perfect symmetry and where the tip atom motif is spread across multiple subunits.
2. **Tip atoms.** For these cases, we evaluate the ability of RFdiffusion3 to scaffold the full symmetrized motif. The tip atoms selected for each copy of the asymmetric unit are the same as in Section 3.5. The total number of residues in the scaffolded motifs ranges from 1 to 7.
3. **ASU size.** For each design, we sample ASU sizes that are equivalent (rounded to the nearest 5 residues) to the length of the native enzyme from which the motif is taken. For examples that do not fit in memory, the length is 290 residues per ASU.

For each motif, we generate 100 designs with RFdiffusion3, sample 8 sequences per backbone with LigandMPNN, and then fold the entire complex across five AlphaFold3 models (one model seed). A successful design must have at least 1 sequence that refolds with a complex C$\alpha$ RMSD of $\leq 2.0$ Å and without any ligand clashes (defined as in Section 3.5). Furthermore, the design must satisfy the motif-recapitulation criteria given in Section 3.5 ; however, we compute the motif RMSD with respect to only the ASU, and the fidelity to the design model with the complex C$\alpha$ RMSD.

##### 3.8 Benchmarking Additional Conditioning

**Hydrogen bond conditioning.** Hydrogen bond conditioning analysis (Fig. 2c) was done for binder design against the target OQO (PDB ID 7V11) and the DNA structure from the paired region of the biological assembly 1 of PDB ID 7RTE. For OQO, three independent conditioning cases were evaluated, each of which annotated a single atom for hydrogen bond conditioning (N5 as donor, O25 as acceptor, and N36 as acceptor). For DNA, three independent cases were evaluated. Each time, a set of two atoms on the major groove side of a base was annotated: A12 (adenine: N7 acceptor, N6 donor), B23 (guanine: N7 acceptor, O6 acceptor), and B24 (guanine: N7 acceptor, O6 acceptor). These atoms are known to be important for DNA base recognition [37]). To explore the value of classifier-free guidance in this setting, each of the above tasks was run both with and without classifier-free guidance (CFG scale 1.5). To isolate the effect of the hydrogen bond conditioning, both examples were also run without any hydrogen bond condtioning. For each design case 100 RFdiffusion3 backbones were generated (`step_scale` 1.5, lengths 50-110 for DNA, 120-150 for OQO). For DNA, a COM specification was set in the major groove region (28.134, -11.888, -22.271). We compute the percentage of specified atomic conditions satisfied and report the average values in (Fig. 2c).

**RASA conditioning** For RASA conditioning analysis (Fig. 2d), the small molecule IAI was chosen (PDB ID: 5SDV). Two conditions were evaluated: fully buried and partially buried. For the partially buried case the following atoms were annotated to be exposed: C22, C23, C25, C24, C21, C20, N13, C15, C16, N14, C19, C11, N12, C18, C17. All other atoms are annotated as 0. All atoms were annotated as buried for the fully buried case. For each case, 100 RFdiffusion backbones were generated, with `step_scale` 1.5 and `CFG_scale` 2. RASA values were computed for each atom of IAI for all designs. Distribution of average RASA values are presented for buried and exposed atoms separately.

#### 4 *In vitro* Evaluation

##### 4.1 Cysteine Hydrolase

Cysteine hydrolases catalyze ester and amide hydrolysis through a double-displacement mechanism utilizing a cysteine thiol as the nucleophile. The reaction proceeds via two tetrahedral intermediates stabilized by an oxyanion hole. In the first step, the deprotonated cysteine thiolate attacks the substrate carbonyl carbon to form a tetrahedral intermediate (TI1), which collapses to yield an acyl–enzyme intermediate (AEI) with release of the alcohol leaving group. The AEI is subsequently attacked by an activated water molecule, forming a second tetrahedral intermediate (TI2) that resolves to regenerate the free enzyme and release the acid product. In natural cysteine hydrolases, a Cys–His–Asp/Asn catalytic triad provides the nucleophile and acid/base chemistry, while backbone and side-chain donors stabilize the negatively charged intermediates. Efficient catalysis requires sub-angstrom precision in the arrangement of these functional groups, making cysteine hydrolases a stringent benchmark for structure-based enzyme design. In RFdiffusion2 [3], the authors demonstrated that active cysteine hydrolases acting on 4-methylumbelliferyl (4MU) ester substrates could be generated *de novo*, establishing this class as a challenging but tractable experimental benchmark. Here, we sought to test whether advances in backbone generation from RFdiffusion3, together with an updated downstream sequence design pipeline using LigandMPNN [26] and AlphaFold3 (AF3) [11] could further improve experimental success in this system.

Using the pipeline described in the following section, we generated 190 unique sequences from 27 distinct scaffolds generated using RFD3 and ordered them as eBlocks (IDT). All designs were screened directly in IVTT reactions using 4-methylumbelliferyl phenyl acetate (4MU–PhAc) as substrate (Fig. S8a). Because this substrate is highly activated, background hydrolysis from the IVTT mixture itself produces a measurable signal; thus, only designs that substantially exceed this background can be considered active. To establish a quantitative threshold, we defined hits as designs with steady-state turnover (5–10 min window) greater than three standard deviations above the mean of six background replicates. By this metric, 73 of 190 designs (38%) exhibited detectable activity. To further identify multi-turnover enzymes, we required that the background-subtracted signal exceed 10% of the maximum fluorescence corresponding to complete substrate conversion (dashed black line, Fig. S8a). This threshold is stringent: given that mScarlet fluorescence across IVTT reactions corresponds to enzyme concentrations below 1 $\mu$M, designs surpassing 10% conversion must have executed multiple catalytic turnovers rather than single-turnover events. By this criterion, 35 of 190 designs (18%) qualified as multi-turnover hits. This represents a substantial increase in success over prior de novo cysteine or serine hydrolase campaigns [3, 68], where one-shot design typically yielded 1–2% multi-turnover success. These results highlight that RFdiffusion3, especially when coupled to iterative LigandMPNN–AF3 sequence refinement, significantly improves the likelihood of producing catalytically competent enzymes. Normalized steady-state rates (background-subtracted and scaled by mScarlet fluorescence to account for expression differences) for all designs are shown in Fig. S8b. Designs are named by their parent scaffold family (alphabetical) and indexed numerically within each family. Of the 27 initial scaffolds, 10 produced at least one multi-turnover hit. Scaffold C was particularly productive: 6 of its 8 sequences were multi-turnover, including the overall best design, C6. The activity of C6 was reproducible following expression and purification from *E. coli*, enabling detailed kinetic analysis (Fig. S8 and Fig. 4cd). The enzyme displayed a catalytic efficiency of $k_{\mathrm{cat}}/K_M = 3.6 \times 10^3\,\mathrm{M}^{-1}\,\mathrm{s}^{-1}$, more than an order of magnitude higher than the best cysteine hydrolase (*Eta1*) from RFdiffusion2 [3]. Moreover, unlike *Eta1*, which exhibited burst-phase kinetics indicative of a rate-limiting deacylation step, C6 showed clean steady-state behavior, suggesting improved active-site pre-organization in this design.

AF3 structural predictions for C6 provide a rationale for its activity. The binding pocket is highly shape-complementary to the modeled tetrahedral intermediate (TI1) and conforms well to the RASA conditioning, burying the acyl moiety while leaving the 4MU group solvent-exposed (Fig. S8e). This complementarity is mechanistically significant: enzymes accelerate reactions by preferentially stabilizing the transition state, and close geometric matching to the tetrahedral intermediate enforces the correct positioning of the scissile bond relative to the catalytic cysteine while disfavoring non-productive poses. Moreover, the catalytic residues are pre-organized with sub-angstrom precision, with the Cys–His–Asp triad and oxyanion-stabilizing groups maintaining the intended hydrogen-bonding geometry (Fig. S8d). Importantly, the Asp general base is buttressed by dual hydrogen bonds from two arginines, whose guanidinium groups are themselves supported by an Asp and Thr on nearby helices. This nested network locks both the Asp and His in place. Separately, the helix–turn–loop motif surrounding the nucleophile is reinforced by an additional hydrogen-bond network to other regions of the protein, which helps stabilize the loop conformation and maintain the backbone oxyanion hole in the correct orientation. Together, these features suggest that RFdiffusion3 can generate backbone scaffolds that can support intricate hydrogen-bond networks and precise active-site preorganization—hallmarks of efficient catalysis.

###### 4.1.1 Design Pipeline

To construct the theozyme, we utilized the crystal structure of the Ulp1 cysteine hydrolase (PDB ID: `1EUV`) which features the canonical Cys-His-Asp triad to construct the catalytic motif by modeling the first tetrahedral intermediate with a 4-methylumbelliferyl phenyl acetate substrate. A tetrahedral conformer was generated at the scissile carbonyl carbon using RDKit and oriented according to established geometric parameters for nucleophilic attack ($\alpha_{\text{cat}} \sim 107°$) and oxyanion stabilization ($\chi_{\text{oxy}} \sim 30°$), while also ensuring proper hydrogen-bonding distances and orientations to the backbone amide nitrogen of Cys25 and the side-chain nitrogen of Gln19. The motif incorporated fixed atomic coordinates from His159 (C$\beta$ and all atoms of the imidazole ring), Asp175 (C$\beta$, C$\gamma$, C$\delta$, and both carboxylate oxygens), Gln19 ($N_{\varepsilon 2}$, $O_{\varepsilon 1}$, C$\delta$, and C$\gamma$), and all heavy atoms of Cys25, together with the complete 4MU–phenyl acetate substrate. To position the nucleophile at the N-terminus of a helix and enhance oxyanion stabilization through the helix dipole, we additionally fixed the backbone $N$, $C\alpha$, $C$ and $O$ atoms of Ser24, as well as the $N$, $C\alpha$, $C$, and $O$ atoms of Trp26.

The resulting theozyme represents a minor refinement of the motif used in RFdiffusion2, incorporating the Ulp1 triad and phenyl acetate substrate analog, and was applied as unindexed motif conditioning input for RFdiffusion3. In addition to motif conditioning, we applied atom-wise relative solvent accessibility (RASA) conditioning to enforce a catalytically sensible exposure pattern for the bound substrate: atoms belonging to the 4MU leaving group were targeted to remain solvent-exposed, whereas atoms in the phenyl-acetate acyl moiety were targeted to be solvent-buried. This differential accessibility promotes a pocket geometry that seats the scissile carbonyl deep enough to support nucleophilic approach by Cys S$\gamma$, while orienting the leaving group toward bulk solvent to facilitate egress and to avoid non-productive burial of the fluorophore, as well as preserving a solvent ingress pathway for water recruitment and attack during acyl–enzyme hydrolysis (AEI $\rightarrow$ TI2). RASA targets were specified at the atom level for the substrate during backbone generation and subsequently enforced as acceptance criteria during backbone filtering. Using this setup, we generated a total of 130,000 scaffolds with RFdiffusion3. We then applied a two-part coarse filter: (i) a local topology check to verify formation of the intended helix–turn–loop architecture around the active site (nucleophile anchored at the N-terminus of an $\alpha$-helix), and (ii) compliance with the prescribed RASA conditioning for the substrate atoms. Approximately

20% of scaffolds met both criteria; thus, ∼26,000 backbones were advanced to sequence design.

Sequence design proceeded through three rounds of LigandMPNN sampling interleaved with AlphaFold3 (AF3) evaluation of the first tetrahedral intermediate (TI1). We employed this iterative sequence–structure refinement loop to drive the system toward a self-consistent sequence–structure pair under the catalytic motif. At each round, AlphaFold3 provides a locally relaxed structural hypothesis for TI1; conditioning LigandMPNN on that hypothesis focuses sequence sampling on variants that stabilize the predicted geometry. Tightening the filters across rounds acts as a continuation scheme, progressively enriching for designs that satisfy catalytic distances/angles with high model confidence. For AF3 evaluation, the first tetrahedral intermediate (TI1) was encoded explicitly by representing the acylation state as a custom non-canonical thioacylated cysteine, ensuring that the nucleophile–carbonyl tetrahedral center was present during prediction. For each backbone in each round, we sampled 10 sequences with LigandMPNN and generated an AF3 ensemble of five TI1 predictions (5 diffusion samples). Round 1 filtering required an ensemble-average motif heavy-atom RMSD $< 1.25$ Å (computed over all theozyme atoms and, from the substrate, only the scissile carbonyl carbon, the carbonyl oxygen, and the leaving-group oxygen) and ensemble-average pTM $> 0.7$; only AF3 predictions with the correct stereochemistry at the tetrahedral center were retained, and designs with no correctly chiral member were discarded. Under these criteria, 5,492 sequences passed, originating from 2,586 distinct scaffolds. The passing AF3 TI1 models were recycled as conditioning structures for LigandMPNN, and Round 2 sampled 10 sequences per design. At this stage we applied stricter, geometry-forward filters to enrich for active-site preorganization: ensemble-average motif RMSD $< 1.0$ Å and pTM $> 0.8$, together with distance constraints (means over the five-member AF3 ensemble) capturing the catalytic arrangement as in prior work—Cys $S_\gamma$–His $N_{\delta 1}$ (nucleophile–base) $\leq 4.0$ Å, His $N_{\delta 1}$–leaving-group O (general acid/base contact) $\leq 4.0$ Å, and carbonyl O–backbone N of the nucleophile (oxyanion–backbone H-bond) $\leq 3.5$ Å. This stringent filter yielded 406 sequences spanning 150 scaffolds. For Round 3, each of these 406 designs was used to condition LigandMPNN to sample 100 sequences, which were evaluated with AF3 (five-member ensembles) and filtered with tightened cutoffs: Cys $S_\gamma$–His $N_{\delta 1} \leq 4.0$ Å, His $N_{\delta 1}$–leaving-group O $\leq 3.8$ Å, carbonyl O–backbone N (nucleophile) $\leq 3.5$ Å, His–Asp (general base–acid partner) $\leq 3.5$ Å, and carbonyl O–side-chain donor (Gln $N_{\varepsilon 2}$ in the oxyanion network) $\leq 3.8$ Å; in addition, we required per-residue pLDDT $> 80$ for each catalytic residue (Cys, His, Asp, and the oxyanion side-chain donor). After Round 3, we applied the Round 3 geometric criteria uniformly to all designs produced across rounds and advanced those meeting the thresholds, yielding 2,548 sequences from 50 unique scaffolds.

Consolidation began with AF3 predictions of the enzyme–substrate (holo/ES) complex to confirm that the ligand binds in the intended pocket and in a pose consistent with the TI1 geometry. We required (i) low cross-interface uncertainty (minimum chain-pair PAE $< 5$ Å), (ii) retention of the TI1 substrate pose (RMSD $< 3$ Å computed over the scissile carbonyl C, the carbonyl O, and the leaving-group O after alignment to the TI1 motif), and (iii) catalytically sensible ES contacts: His $N_{\delta 1}$ to leaving-group O $\leq 6$ Å and carbonyl O to the nucleophile's backbone amide N $\leq 6$ Å, capturing general acid/base proximity and oxyanion preorganization, respectively. Designs failing any ES gate were discarded, leaving sequences from 27 scaffolds. For the surviving set we predicted AEI, TI1, and TI2 and assigned each design a composite score built from per-state geometric and confidence terms. For holo/AEI/TI1/TI2 we aggregated: motif heavy-atom RMSD (with $\leq$ 1 Å targets for transition-state analogs), catalytic distances (Cys $S_\gamma$–His $N_{\delta 1}$, His–Asp, His $N_{\delta 1}$–leaving-group O, carbonyl O–Cys backbone N, and carbonyl O–Gln $N_{\varepsilon 2}$), AF3 interface uncertainty for bound states (minimum chain-pair PAE target of 2 Å), retention of the TI1 substrate pose (RMSD $\leq$ 2.5 Å computed over the scissile carbonyl C, the carbonyl O, and the leaving-group O after alignment to the TI1 motif), and confidence thresholds (per-residue pLDDT $\geq$ 90 for catalytic residues and chain-level pLDDT $\geq$ 90 for the protein). Each metric contributed an equal-weight, piecewise score that rewarded meeting or surpassing its target and imposed an exponential penalty otherwise; the composite was the arithmetic mean of these scores. We then ranked sequences within each backbone family by the composite and selected a variable top-$k$ per family (2–10, typically ∼3) to allocate more slots to highly designable backbones while maintaining diversity, assembling 96 designs. For each of these 96 passing designs, we performed a redesign pass (100 LigandMPNN samples per design, AF3 evaluation across all four states with identical composite scoring) and chose the single top-scoring sequence per design, yielding a second set of 96 designs. Two sequences failed DNA synthesis quality control, and the remaining 190 were ordered and advanced to experimental testing.

###### 4.1.2 Experimental Methods

###### Initial Activity Screen

IVTT templates were generated by Golden Gate assembly followed by PCR amplification. Screening was performed in 384-well plates at a final reaction volume of 6 $\mu$L per well. Each well contained 600 nL PUREfrex 2.1 IVTT mix; the remaining 5.4 $\mu$L comprised assay buffer (100 mM HEPES, 50 mM NaCl, pH 7.4, 5% DMSO) and substrate, adjusted such that the *final* reaction contained 100 $\mu$M 4-methylumbelliferyl phenyl acetate (4MU–PhAc). For every design, IVTT reactions, no-enzyme IVTT controls, and buffer-only controls were each run in technical triplicate. Product formation (4-methylumbelliferone, 4MU) was monitored at 25 °C on a Neo2 plate reader (BioTek) using 365/445 nm excitation/emission. Background signals from matched controls were subtracted, and a 4MU standard curve (identical buffer/DMSO) was used to convert fluorescence to concentration. Designs showing a reproducible increase over background in triplicate were designated as hits. The best performing design was further expression in *E.coli*, purified and further characterized to obtain its Michaelis-Menten parameters.

###### Kinetic Analysis and Hit Validation

**Protein Expression and Purification** Synthetic genes (eBlocks, IDT) were assembled by Golden Gate into the LM627 vector bearing C-terminal SNAC and hexahistidine-tag tags. Plasmids were introduced into *E. coli* BL21(DE3) by heat shock, recovered for 1 h at 37 °C in 100 $\mu$L SOC, and transferred to 2 mL deep-well plates containing 900 $\mu$L LB with 50 $\mu g\,mL^{-1}$ kanamycin. Plates were sealed with breathable film and shaken overnight at 37 °C (1200 rpm). For growing cultures, glycerol stocks were prepared and 500 $\mu$L of the overnight was used to inoculate 50 mL autoinduction medium in 250 mL baffled flasks. Cultures were grown 4–6 h at 37 °C, then shifted to 18 °C for overnight expression. Cell pellets were collected (4000 $g$, 15 min, 4 °C) and resuspended in 25 mL wash buffer (50 mM sodium phosphate, 500 mM NaCl, 40 mM imidazole,

1 mM TCEP, pH 7.4) with lysozyme (1 $mg\,mL^{-1}$) and DNase I (0.1 $mg\,mL^{-1}$). Cells were lysed by sonication on ice (2.5 min, 80% amplitude, 10 s on/10 s off) and clarified (14,000 $g$, 30 min, 4 °C). The supernatant was loaded onto 1 mL Ni-NTA resin equilibrated in wash buffer, washed (3 × 15 mL), and eluted with imidazole (pre-elution 400 $\mu$L; final 1.3 mL; 400 mM imidazole in wash buffer). Eluates were further purified by SEC (Superdex 75 Increase 10/300 GL) into 20 mM HEPES, 50 mM NaCl, 1 mM TCEP (pH 7.4), flash forzen and stored at -80 °Cuntil further use. Molecular mass of the protein was confirmed by LC–MS.

**Michaelis–Menten Kinetics.** Michaelis–Menten measurements were performed at 25 °C in 96-well plates (final volume 60 $\mu$L) using assay buffer (100 mM HEPES, 50 mM NaCl, pH 7.4, 5% DMSO). Each well received 6 $\mu$L purified enzyme, resulting in a final enzyme concentration of 0.8 $\mu$M in the reaction (concentration from $A_{280}$ and the computed extinction coefficient). Substrate dependence was recorded over an eight-point, 2× serial dilution starting at 100 $\mu$M 4MU–PhAc (final), with technical triplicates. Initial rates ( $\leq$10% conversion) were background-subtracted using matched controls, converted to concentration via a 4MU standard curve, and fit by nonlinear least squares to obtain $k_{\text{cat}}$ and $K_M$.

#### 4.2 DNA-binder

For the purposes of a proof-of-principle experiment of DNA-binder generation with RFdiffusion3, we used a fine-tuned version of RFdiffusion3 that was trained on a higher percentage of the DNA distillation and DNA interfaces datasets in Section 1.10. AlphaFold3 (AF3) [11] prediction of a random target DNA sequence (random 10 base pairs flanked by CG on either end) was taken as the input DNA motif. Multiple Center of mass (COM) specifications were sampled for running diffusion: one per each 8 base-pair (bp) window. For a given 8-bp window, a COM specification was computed as described in supplementary Section 3.3. For each COM case, 50 protein backbones of length 50-80 were generated with the fine-tuned RFdiffusion3. LigandMPNN [26] was used to design 8 sequences per backbone. These designs were folded with AF3 and designs with less than 3 Å DNA-aligned protein RMSD were chosen to take into a resampling step, which we found to improve diversity and agreement with AF3.

In the resampling step, for each candidate design we selected an indexed protein motif by taking the 10-residue contiguous region which made the most contacts with hydrogen bond donor/acceptor atoms in the DNA major groove. The fine-tuned RFdiffusion3 was used to motif-scaffold this protein motif and the DNA to generate 50 backbones per candidate: 25 designs with 15-35 diffused amino acids on either side of the indexed protein motif and 25 designs with 25-50 diffused residues on either side. LigandMPNN was used to design 8 sequences per backbone. Designs with less than 3 Å AF3 DNA-aligned protein RMSD, higher than 0.8 iPTM and length less than 87 were chosen. These high confidence designs were ordered as synthetic genes (Gene Fragments, Twist Bioscience) and transformed into Saccharomyces cerevisiae strain EBY100 using 2–3 µg of pETcon vector (pETcon3), for yeast surface display. Yeast cultures were grown in C-Trp-Ura medium with 2% glucose and induced in SGCAA medium supplemented with 0.2% glucose at a density of $1 \times 10^7$ cells/ml for 16–24 h at 30 °C. Cells were washed with PBSF (PBS with 1% BSA) and labeled with biotinylated dsDNA target oligos, anti-c-Myc FITC (Miltenyi Biotech), and streptavidin–PE (SAPE, ThermoFisher) with avidity. One out of the 5 designs (named DBRFD3) ordered for target `CGAGAACATAGTCG` was found to be binding to its cognate DNA target, at 1 $\mu$M concentration. Structural analysis of the AF3 prediction of DBRFD3 reveals a recognition mechanism driven through extensive hydrogen bonding, polar, and non-polar interactions with the DNA major groove and backbone Fig. S7.

To further characterize this hit, we performed a three-replicate titration series using biotinylated dsDNA target oligos ranging from ∼10 nM to 30 µM under no avidity conditions. Flow cytometry data were analyzed using custom Python code in combination with the CytoFlow package: expression-positive yeast cells were gated using a Gaussian Mixture Model, and for each sample the median fluorescence intensity in the SAPE (binding) channel was normalized to the FITC (expression) channel to yield a binding-to-expression ratio. Fitting of the non-avidity titration data to a four-parameter logistic regression (with top and bottom constrained to 100% and 0%) yielded an apparent EC50 of $5.89 \pm 2.15$ µM, estimated based on three replicates. These results demonstrates that DBRFD3 binds the target DNA sequence with low micromolar affinity in the absence of avidity effects.

While some native transcription factors (TFs) such as NF-$\kappa$B or homeodomains can bind with sub-nanomolar strength [69, 70], others, including MYC:MAX (∼0.5 µM) and FoxA1 (∼1 µM), exhibit affinities in the high-nanomolar to micromolar range [71, 72]. This information puts affinity of DBRFD3 into context. Future optimization could leverage targeted mutagenesis, which has successfully increased the affinity of native binders such as homeodomains and zinc fingers [73, 74], and represents a natural next step for optimizing de novo designs. Follow up studies will explore broader scale design of DNA binding proteins with RFdiffusion3.

[4] Martin Pacesa, Lennart Nickel, Christian Schellhaas, Joseph Schmidt, Ekaterina Pyatova, Lucas Kissling, Patrick Barendse, Jagrity Choudhury, Srajan Kapoor, Ana Alcaraz-Serna, Yehlin Cho, Kourosh H. Ghamary, Laura Vinué, Brahm J. Yachnin, Andrew M. Wollacott, Stephen Buckley, Adrie H. Westphal, Simon Lindhoud, Sandrine Georgeon, Casper A. Goverde, Georgios N. Hatzopoulos, Pierre Gönczy, Yannick D. Muller, Gerald Schwank, Daan C. Swarts, Alex J. Vecchio, Bernard L. Schneider, Sergey Ovchinnikov, and Bruno E. Correia. One-shot design of functional protein binders with BindCraft. *Nature*, 2025. URL https://doi.org/10.1038/s41586-025-09429-6.

[5] Jason Yim, Andrew Campbell, Emile Mathieu, Andrew Y. K. Foong, Michael Gastegger, Jose Jimenez-Luna, Sarah Lewis, Victor Garcia Satorras, Bastiaan S. Veeling, Frank Noe, Regina Barzilay, and Tommi Jaakkola. Improved motif-scaffolding with SE(3) flow matching. In *ICLR 2024 Workshop on Generative and Experimental Perspectives for Biomolecular Design*, 2024. URL https://openreview.net/forum?id=96JxNcA9r6.

[6] Anthony Marchand, Stephen Buckley, Arne Schneuing, Martin Pacesa, Maddalena Elia, Pablo Gainza, Evgenia Elizarova, Rebecca M. Neeser, Pao-Wan Lee, Luc Reymond, Yangyang Miao, Leo Scheller, Sandrine Georgeon, Joseph Schmidt, Philippe Schwaller, Sebastian J. Maerkl, Michael Bronstein, and Bruno E. Correia. Targeting protein–ligand neosurfaces with a generalizable deep learning tool. *Nature*, 639(8054):522–531, 2025. doi: 10.1038/s41586-024-08435-4.

[7] John B. Ingraham, Max Baranov, Zak Costello, Karl W. Barber, Wujie Wang, Ahmed Ismail, Vincent Frappier, Dana M. Lord, Christopher Ng-Thow-Hing, Erik R. Van Vlack, Shan Tie, Vincent Xue, Sarah C. Cowles, Alan Leung, João V. Rodrigues, Claudio L. Morales-Perez, Alex M. Ayoub, Robin Green, Katherine Puentes, Frank Oplinger, Nishant V. Panwar, Fritz Obermeyer, Adam R. Root, Andrew L. Beam, Frank J. Poelwijk, and Gevorg Grigoryan. Illuminating Protein Space with a Programmable Generative Model. *Nature*, 623(7989): 1070–1078, 2023. URL https://doi.org/10.1038/s41586-023-06728-8.

[8] Vinicius Zambaldi, David La, Alexander E Chu, Harshnira Patani, Amy E Danson, Tristan OC Kwan, Thomas Frerix, Rosalia G Schneider, David Saxton, Ashok Thillaisundaram,

et al. De novo design of high-affinity protein binders with AlphaProteo. *arXiv preprint arXiv:2409.08022*, 2024.

[9] Jue Wang, Sidney Lisanza, David Juergens, Doug Tischer, Joseph L. Watson, Karla M. Castro, Robert Ragotte, Amijai Saragovi, Lukas F. Milles, Minkyung Baek, Ivan Anishchenko, Wei Yang, Derrick R. Hicks, Marc Expòsit, Thomas Schlichthaerle, Jung-Ho Chun, Justas Dauparas, Nathaniel Bennett, Basile I. M. Wicky, Andrew Muenks, Frank DiMaio, Bruno Correia, Sergey Ovchinnikov, and David Baker. Scaffolding protein functional sites using deep learning. *Science*, 377(6604):387–394, 2022. doi: 10.1126/science.abn2100. URL https://www.science.org/doi/abs/10.1126/science.abn2100.

[10] Chai Discovery Team, Jacques Boitreaud, Jack Dent, Danny Geisz, Matthew McPartlon, Joshua Meier, Zhuoran Qiao, Alex Rogozhnikov, Nathan Rollins, Paul Wollenhaupt, and Kevin Wu. Zero-shot antibody design in a 24-well plate. *bioRxiv*, 2025. doi: 10.1101/2025.07.05.663018. URL https://www.biorxiv.org/content/early/2025/07/06/2025.07.05.663018.

[11] Josh Abramson, Jonas Adler, Jack Dunger, Richard Evans, Tim Green, Alexander Pritzel, Olaf Ronneberger, Lindsay Willmore, Andrew J Ballard, Joshua Bambrick, et al. Accurate structure prediction of biomolecular interactions with AlphaFold 3. *Nature*, pages 1–3, 2024.

[12] Tomas Geffner, Kieran Didi, Zuobai Zhang, Danny Reidenbach, Zhonglin Cao, Jason Yim, Mario Geiger, Christian Dallago, Emine Kucukbenli, Arash Vahdat, et al. Proteina: Scaling flow-based protein structure generative models. *arXiv preprint arXiv:2503.00710*, 2025.

[13] Tomas Geffner, Kieran Didi, Zhonglin Cao, Danny Reidenbach, Zuobai Zhang, Christian Dallago, Emine Kucukbenli, Karsten Kreis, and Arash Vahdat. La-Proteina: Atomistic Protein Generation via Partially Latent Flow Matching, 2025. arXiv preprint.

[14] Nathaniel Corley, Simon Mathis, Rohith Krishna, Magnus S Bauer, Tuscan R Thompson, Woody Ahern, Maxwell W Kazman, Rafael I Brent, Kieran Didi, Andrew Kubaney, Lilian McHugh, Arnav Nagle, Andrew Favor, Meghana Kshirsagar, Pascal Sturmfels, Yanjing Li, Jasper Butcher, Bo Qiang, Lars L Schaaf, Raktim Mitra, Katelyn Campbell, Odin Zhang, Roni Weissman, Ian R Humphreys, Qian Cong, Jonathan Funk, Shreyash Sonthalia, Pietro Liò, David Baker, and Frank DiMaio. Accelerating biomolecular modeling with AtomWorks and RF3. Preprint available at bioRxiv, 2025.

[20] Jonathan Ho and Tim Salimans. Classifier-free diffusion guidance. *arXiv preprint arXiv:2207.12598*, 2022.

[21] Helen M Berman, John Westbrook, Zukang Feng, Gary Gilliland, Talapady N Bhat, Helge Weissig, Ilya N Shindyalov, and Philip E Bourne. The protein data bank. *Nucleic acids research*, 28(1):235–242, 2000.

[22] wwPDB consortium. Protein Data Bank: the single global archive for 3D macromolecular structure data. *Nucleic acids research*, 47(D1):D520–D528, 2019.

[28] Longxing Cao, Brian Coventry, Inna Goreshnik, Buwei Huang, William Sheffler, Joon Sung Park, Kevin M Jude, Iva Marković, Rameshwar U Kadam, Koen H G Verschueren, Kenneth Verstraete, Scott Thomas Russell Walsh, Nathaniel Bennett, Ashish Phal, Aerin Yang, Lisa Kozodoy, Michelle DeWitt, Lora Picton, Lauren Miller, Eva-Maria Strauch, Nicholas D DeBouver, Allison Pires, Asim K Bera, Samer Halabiya, Bradley Hammerson, Wei Yang, Steffen Bernard, Lance Stewart, Ian A Wilson, Hannele Ruohola-Baker, Joseph Schlessinger, Sangwon Lee, Savvas N Savvides, K Christopher Garcia, and David Baker. Design of protein-binding proteins from the target structure alone. *Nature*, 605(7910):551–560, May 2022.

[38] Nathan M Ennist, Shunzhi Wang, Madison A Kennedy, Mariano Curti, George A Sutherland, Cvetelin Vasilev, Rachel L Redler, Valentin Maffeis, Saeed Shareef, Anthony V Sica, Ash Sueh Hua, Arundhati P Deshmukh, Adam P Moyer, Derrick R Hicks, Avi Z Swartz, Ralph A Cacho, Naia Novy, Asim K Bera, Alex Kang, Banumathi Sankaran, Matthew P Johnson, Amala Phadkule, Mike Reppert, Damian Ekiert, Gira Bhabha, Lance Stewart, Justin R Caram, Barry L Stoddard, Elisabet Romero, C Neil Hunter, and David Baker. De novo design of proteins housing excitonically coupled chlorophyll special pairs. *Nat. Chem. Biol.*, 20(7): 906–915, 2024.

[47] Anna Lauko, Samuel J. Pellock, Kiera H. Sumida, Ivan Anishchenko, David Juergens, Woody Ahern, Jihun Jeung, Alexander F. Shida, Andrew Hunt, Indrek Kalvet, Christoffer Norn, Ian R. Humphreys, Cooper Jamieson, Rohith Krishna, Yakov Kipnis, Alex Kang, Evans

Brackenbrough, Asim K. Bera, Banumathi Sankaran, K. N. Houk, and David Baker. Computational design of serine hydrolases. *Science*, 388(6744):eadu2454, 2025. doi: 10.1126/science.adu2454. URL https://www.science.org/doi/abs/10.1126/science.adu2454.

[56] Minkyung Baek, Frank DiMaio, Ivan Anishchenko, Justas Dauparas, Sergey Ovchinnikov, Gyu Rie Lee, Jue Wang, Qian Cong, Lisa N Kinch, R. Dustin Schaeffer, Claudia Millán, Hahnbeom Park, Carlos Adams, Craig R Glassman, Alexander DeGiovanni, Jose H Pereira, Andrew V Rodrigues, Amelie A van Dijk, Andrea C Ebrecht, D. J. Opperman, Tobias Sagmeister, Christoph Buhlheller, Tea Pavkov-Keller, Manoj K Rathinaswamy, Ujwala Dalwadi, Christopher K Yip, John E Burke, K. Christopher Garcia, Nick V Grishin, Paul D Adams, Randy J Read, and David Baker. Accurate prediction of protein structures and interactions using a three-track neural network. *Science*, 373(6557):871–876, 2021. doi: 10.1126/science.abj8754. URL https://www.science.org/doi/10.1126/science.abj8754.

[57] Jinsen Li, Tsu-Pei Chiu, and Remo Rohs. Deep DNAshape: Predicting DNA shape considering extended flanking regions using a deep learning method. *bioRxiv : the preprint server for biology*, 10 2023. doi: 10.1101/2023.10.22.563383.

[58] Xiang-Jun Lu and Wilma K Olson. 3DNA: a software package for the analysis, rebuilding and visualization of three-dimensional nucleic acid structures. *Nucleic acids research*, 31(17): 5108–5121, 2003.

[59] Raktim Mitra, Ari S Cohen, Jared M Sagendorf, Helen M Berman, and Remo Rohs. DNAproDB: an updated database for the automated and interactive analysis of protein–DNA complexes. *Nucleic acids research*, 53(D1):D396–D402, 2025.

[60] Tero Karras, Miika Aittala, Timo Aila, and Samuli Laine. Elucidating the Design Space of Diffusion-Based Generative Models. In S. Koyejo, S. Mohamed, A. Agarwal, D. Belgrave, K. Cho, and A. Oh, editors, *Advances in Neural Information Processing Systems*, volume 35, pages 26565–26577. Curran Associates, Inc., 2022.

[61] Ian K McDonald and Janet M Thornton. Satisfying hydrogen bonding potential in proteins. *Journal of molecular biology*, 238(5):777–793, 1994.

[62] Andrew Shrake and John A Rupley. Environment and exposure to solvent of protein atoms. Lysozyme and insulin. *Journal of molecular biology*, 79(2):351–371, 1973.

[63] Patrick Kunzmann and Kay Hamacher. Biotite: a unifying open source computational biology framework in Python. *BMC bioinformatics*, 19(1):346, 2018.

[64] Barbara Zdrazil, Eloy Felix, Fiona Hunter, Emma J Manners, James Blackshaw, Sybilla Corbett, Marleen de Veij, Harris Ioannidis, David Mendez Lopez, Juan F Mosquera, Maria Paula Magarinos, Nicolas Bosc, Ricardo Arcila, Tevfik Kizilören, Anna Gaulton, A Patrícia Bento, Melissa F Adasme, Peter Monecke, Gregory A Landrum, and Andrew R Leach. The ChEMBL Database in 2023: a drug discovery platform spanning multiple bioactivity data types and time periods. *Nucleic Acids Res.*, 52(D1):D1180–D1192, 2024.
